# Cryo-electron tomography of enterovirus cell entry and endosome escape

**DOI:** 10.1101/2023.07.12.548671

**Authors:** Aygul Ishemgulova, Liya Mukhamedova, Zuzana Trebichalská, Veronika Rájecká, Pavel Payne, Lenka Šmerdová, Jana Moravcová, Dominik Hrebík, David Buchta, Karel Škubník, Tibor Füzik, Štěpánka Vaňáčová, Jiří Nováček, Pavel Plevka

**Author notes:** These authors contributed equally (Aygul Ishemgulova, Liya Mukhamedova, Zuzana Trebichalská).

## Abstract

Enveloped viruses deliver their genomes into the cell cytoplasm by membrane fusion; in contrast, membrane penetration by non-enveloped viruses is more diverse and less well understood. Enteroviruses, one of the largest groups of non-enveloped viruses, cause diseases ranging from the common cold to life-threatening encephalitis. To initiate infection, most enteroviruses enter cells by endocytosis. However, how enterovirus particles or RNA genomes cross the endosome membrane into the cytoplasm remains unknown. Here we used cryo-electron tomography of infected cells to show that endosomes containing rhinovirus 2, echovirus 18, echovirus 30, or enterovirus 71 deform, rupture, and release their content into the cytoplasm. Blocking endosome acidification with bafilomycin A1 reduced the number of enterovirus particles that released their genomes in endosomes, but did not prevent them from reaching the cytoplasm. Inhibiting N-WASP-mediated post-endocytic membrane remodeling with wiskostatin promoted abortive enterovirus genome release in endosomes. We show that the rupture of endosomes also occurs in uninfected cells. In summary, our results indicate that cellular membrane remodeling disrupts enterovirus-containing endosomes and thus releases the virus genomes and particles into the cytoplasm. Since the studied enteroviruses employ different receptors for cell entry but are all delivered into the cytoplasm by cell-mediated endosome disruption, it is possible that many other enteroviruses utilize endosome rupture to infect cells.

## Introduction

Enteroviruses from the family *Picornaviridae* cause a wide range of diseases, including the common cold, hand-foot-and-mouth disease, encephalitis, and paralysis, with major health, economic, and societal impacts (*1–3*). Virions of enteroviruses are formed of capsids with diameters of 30 nm and ∼7,500 nucleotide-long single-stranded RNA genomes (*4*). Enterovirus capsids are built from VP1, VP2, and VP3 subunits organized with pseudo-T = 3 icosahedral symmetry and minor capsid proteins VP4 symmetrically attached to the inner face of the capsid (*4–6*). To initiate infection, enteroviruses bind to receptors, and most of them enter cells by endocytosis (*7–13*). Endocytosis is an essential cellular process of membrane vesicle traffic, which involves the internalization of segments of the plasma membrane and engulfed extracellular cargo into a cell. There are multiple types of endocytosis that function in parallel and can be broadly divided into (i) macroscale phagocytosis or macro-pinocytosis and (ii) microscale clathrin-, caveolin-, and flotillin-mediated as well as coat-independent endocytosis (*14*). Enteroviruses attach to specific cellular receptors; however, they can utilize various types of endocytosis for cell entry (*7–9, 15*). Rhinovirus A and B serotypes are classified into major and minor groups depending on whether they utilize intercellular adhesion molecule 1 or low-density lipoprotein receptor (LDLR) family members for cell entry (*16–18*). Of the viruses analyzed in this study, rhinovirus 2 binds to LDLRs (*17*). The entry of enterovirus 71 is mediated by P-selectin glycoprotein ligand 1, Scavenger receptor class B member 2, and heparan sulfate (*19–21*). Echoviruses 18 and 30 attach to the neonatal Fc receptor, while echovirus 30 additionally utilizes a decay-accelerating factor (*22, 23*). In endosomes, enteroviruses are exposed to acidic pH (*23–25*). *In vitro* studies have shown that receptor binding and acidic pH induce the transformation of enterovirus virions into activated particles, which are characterized by expanded capsids, reduction of contacts between pentamers of capsid proteins, the release of VP4, externalization of N-termini of VP1, and granular distribution of genomic RNA (*26–31*). Activated particles spontaneously release their genomes to enable the initiation of translation and replication (*29, 30, 32, 33*).

Various mechanisms for enterovirus genome delivery into the cytoplasm have been proposed (*8, 34*). It was suggested that poliovirus, rhinovirus 2, and rhinovirus 16 release their genomes in endosomes and translocate them into the cytoplasm through transmembrane pores formed by VP4 and N-termini of VP1 (*35–39*). Poliovirus particles associate with liposomes decorated with extracellular domains of poliovirus receptors and form umbilical connections to the membrane through which the genomes might be transported (*40–45*). However, diffusion of the ∼7,500-nucleotide-long RNA genome through an opening in a capsid and putative transmembrane pore would be a slow process, which has not been verified *in vivo*. Major group rhinoviruses and rhinoviruses which have adapted to use heparan-sulfate as a receptor were indicated to induce endosome disruption and release their genomes both into the endosomes and cytoplasm (*46, 47*). It has been shown that poliovirus and equine-rhinitis A virus genomes are protected from RNases co-endocytosed with the virus particles (*45*). Despite the previous studies, our understanding of enterovirus genome delivery into the cell cytoplasm is incomplete.

Here we used cryo-electron tomography (cryo-ET) to visualize the cell entry of rhinovirus 2, echovirus 18, echovirus 30, and enterovirus 71 *in situ*. We show that enterovirus-containing endosomes deform, rupture, and release the virus particles into the cytoplasm. Blocking endosome acidification with bafilomycin A1 inhibited enterovirus genome release, but did not prevent endosome disruption. Inhibiting actin-dependent post-endocytic membrane remodeling with wiskostatin resulted in the accumulation of empty enterovirus particles in endosomes. We propose that cellular membrane-remodeling machinery disrupts endosomes and thus enables the diffusion of enterovirus particles and genomes into the cytoplasm.

## Results and Discussion

### Endocytosis of enteroviruses visualized *in situ*

We employed cryo-ET to visualize the entry of rhinovirus 2, echovirus 18, echovirus 30, and enterovirus 71 into fibroblast-like cos-7 cells (Fig. 1, 2, S1-3, Movie S1). These cells are permissive to infection by the studied viruses and form thin electron-transparent lamellipodia, which are suitable for transmission electron microscopy imaging without prior thinning. The more than 250 recorded tomograms were of sufficient quality to enable the identification of genome-containing and empty enterovirus particles, ribosomes, microfilaments, microtubules, and cellular membranes (Fig. 1, Movie S1). Seven to sixty minutes post-infection, enterovirus particles were both in membrane-bound vesicles and cytoplasm, indicating that enterovirus cell entry was a continuous and unsynchronized process. With increasing time post-infection, the fractions of empty particles and particles in the cell cytoplasm increased (Fig. S4). Some of the virus-containing endosomes were partially or fully covered with protein coats; however, we could not identify the type of vesicle coating (Fig. 2ab, S1a, S2ab, S3a). Previous studies demonstrated that enteroviruses utilize clathrin-dependent, caveolin-dependent, and other types of endocytosis for cell entry (*7–9, 15*).

**Fig. 1.**
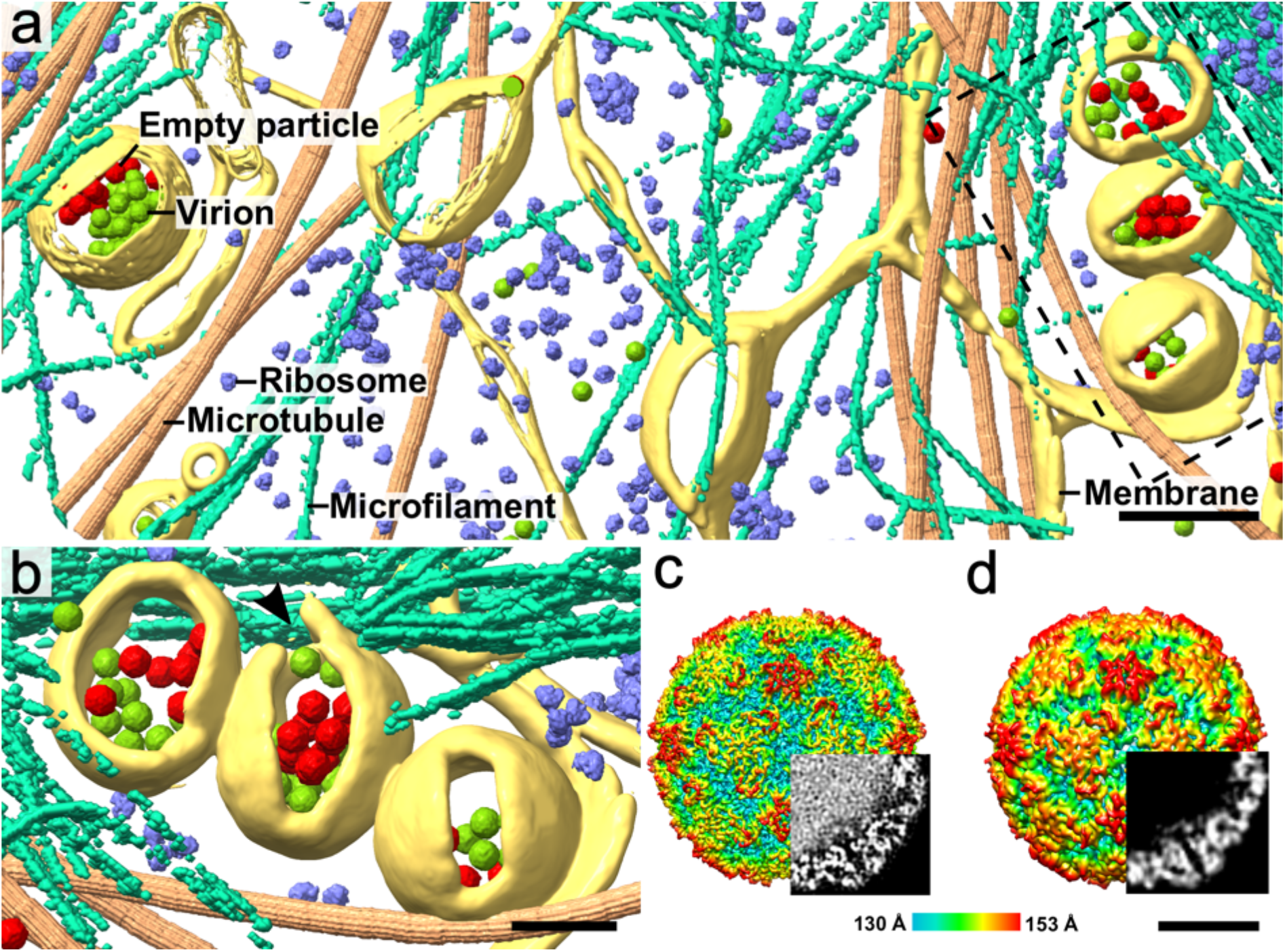
Rhinovirus 2 infection of a cell imaged using cryo-ET. **(a)** Tomographic reconstruction of the cytoplasm of an infected cos-7 cell. Positions of virions, empty particles, ribosomes (EMD-8345) (*84*), and microtubules (EMD-12639) (*85*) were identified using template matching as implemented in EMclarity (*86*), and the corresponding high-resolution structures were placed into the tomogram. Microfilaments and membranes were segmented manually. The dashed rectangle indicates the position of the area shown at higher magnification in panel B. Scale bar 200 nm. **(b)** Detail of endosomes containing virions of rhinovirus 2. The central vesicle is ruptured at the place indicated with a black arrowhead. Scale bar 100 nm. **(cd)** Surface representations of single-particle cryo-EM reconstructions of rhinovirus 2 virion (c) and empty particle (d) calculated from projection images of infected cos-7 cells. The surfaces of the particles are rainbow-colored according to their distance from the particle center. Bottom-right quadrants were replaced with central slices of cryo-EM density distributions within the particles to demonstrate that the virion contains a genome and the empty particle does not. Scale bar 10 nm.

**Fig. 2.**
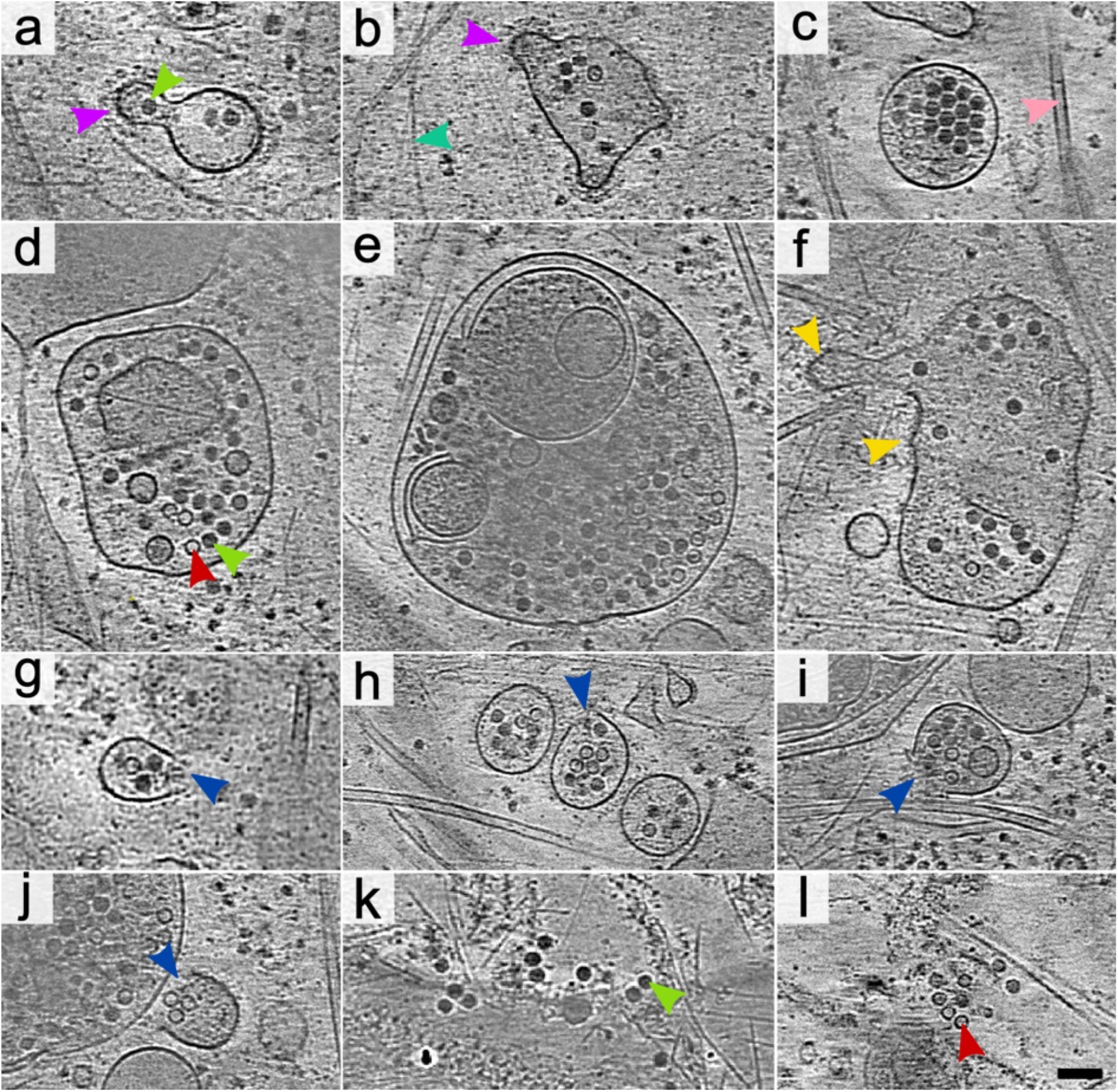
Release of rhinovirus 2 from endosomes into the cytoplasm. The images show 1.66 nm-thick tomographic slices of rhinovirus 2-infected cos-7 cells seven to sixty minutes post-infection. **(ab)** Endosomes with segments of their surface covered with protein coats. Selected coated areas are indicated by magenta arrowheads. A selected virion is indicated by a green arrowhead. A selected actin microfilament is indicated by a cyan arrowhead. **(c)** Single-membrane endosome with a smooth surface containing virions and empty particles of rhinovirus 2. A pink arrowhead indicates a selected microtubule. **(de)** Multilamellar endosomes with a smooth outer membrane containing virions and empty particles. A selected empty particle is indicated by a red arrowhead. **(f)** Endosome with warped membrane forming cone-shaped protrusions. Selected tips of membrane warping are indicated by yellow arrowheads. **(g-j)** Ruptured endosomes release rhinovirus 2 into the cytoplasm. Positions of the openings in endosome membranes are indicated by blue arrows. **(kl)** Virions and empty particles in the cell cytoplasm. Scale bar 100 nm.

### Enterovirus particles do not bind to endosome membranes

The cell entry of enteroviruses is initiated by binding to specific receptors (*4, 17, 19–23*). We observed attachment of the studied enteroviruses to the cytoplasmic membranes of cells, which was presumably mediated by receptor binding (Fig. S5, Movie S2). Some of the particle may have been co-endocytosed as components of solution engulfed into endosomes and, thus, entered cells without binding to receptors. After endocytosis, neither of the studied enteroviruses remained bound to the endosome membrane. Instead, both genome-containing and empty particles were distributed randomly in the endosome lumen (Fig. 2a-f, S1a-c, S2a-c, S3a-c). This confirms previous evidence that rhinovirus 2 detaches from LDLRs soon after endocytosis because of the conformational changes of the receptors induced by low calcium ion concentrations and acidic pH (*48–52*). The mechanisms promoting the detachment of echovirus 18, echovirus 30, and enterovirus 71 from their respective receptors have not been identified. For these viruses, the dissociation is likely triggered by conformational changes to the receptors or virus capsid proteins induced by acidification or changes in ion concentrations in the lumen of maturing endosomes (*31*). Our results indicate that detachment from receptors in endosomes is a common feature of enterovirus cell entry.

Previous studies demonstrated that activated and empty particles of enteroviruses are more hydrophobic than those of virions because of the exposure of N-termini of VP1 subunits at the particle surface (*37, 44, 53, 54*). Furthermore, there is evidence from *in vitro* experiments that activated particles of some enteroviruses bind to membranes and enable translocation of their genomes across the membrane by forming a transmembrane pore composed of N-termini of VP1 and VP4 (*37, 38, 44, 54*). However, we did not observe the binding of the particles of any of the studied enteroviruses to endosome membranes (Fig. 2, S1-3), which suggests that this event may be rare *in vivo* or that the studied enteroviruses employ different mechanism for endosome escape.

### Warping of endosome membranes

Some of the endosomes containing the studied enteroviruses formed cone-shaped protrusions, which we named membrane warping (Fig. 2f, S1c, S2cd, S3c). It should be noted that membranes of a sub-population of endosomes in non-infected cells were also warped, indicating that this is a natural process (Fig. 3a, S6). The observed membrane warping may be related to the previously characterized formation of membrane protrusions, which enable early and sorting endosomes to sort and recycle cargo, receptors, and membranes (*55–57*). The shapes of the membrane cones indicate that they are formed by an external force pulling the membrane away from the endosome center causing membrane deformation (Fig. 2f, 3a, S1c, S2cd, S3c, S6). In some cases, the tips of the membrane cones were attached to actin filaments that may serve as scaffolds for putative complexes, which bind to and pull on the vesicle membranes (Fig. S1c). It was shown that the remodeling of early and sorting endosomes is regulated by the interplay of numerous protein complexes, including endosomal sorting complexes required for transport (ESCRT) (*58*), retromer (*59*), retriever (*60*), nexins (*61*), neural Wiskott–Aldrich syndrome protein (N-WASP) (*62, 63*), Arp 2/3 (*64*), small GTP-binding proteins Rab (*65*), and actin (*66*). It is possible that some of these membrane remodeling complexes contribute to the warping and disruption of enterovirus-containing vesicles, as is discussed below.

**Fig. 3.**
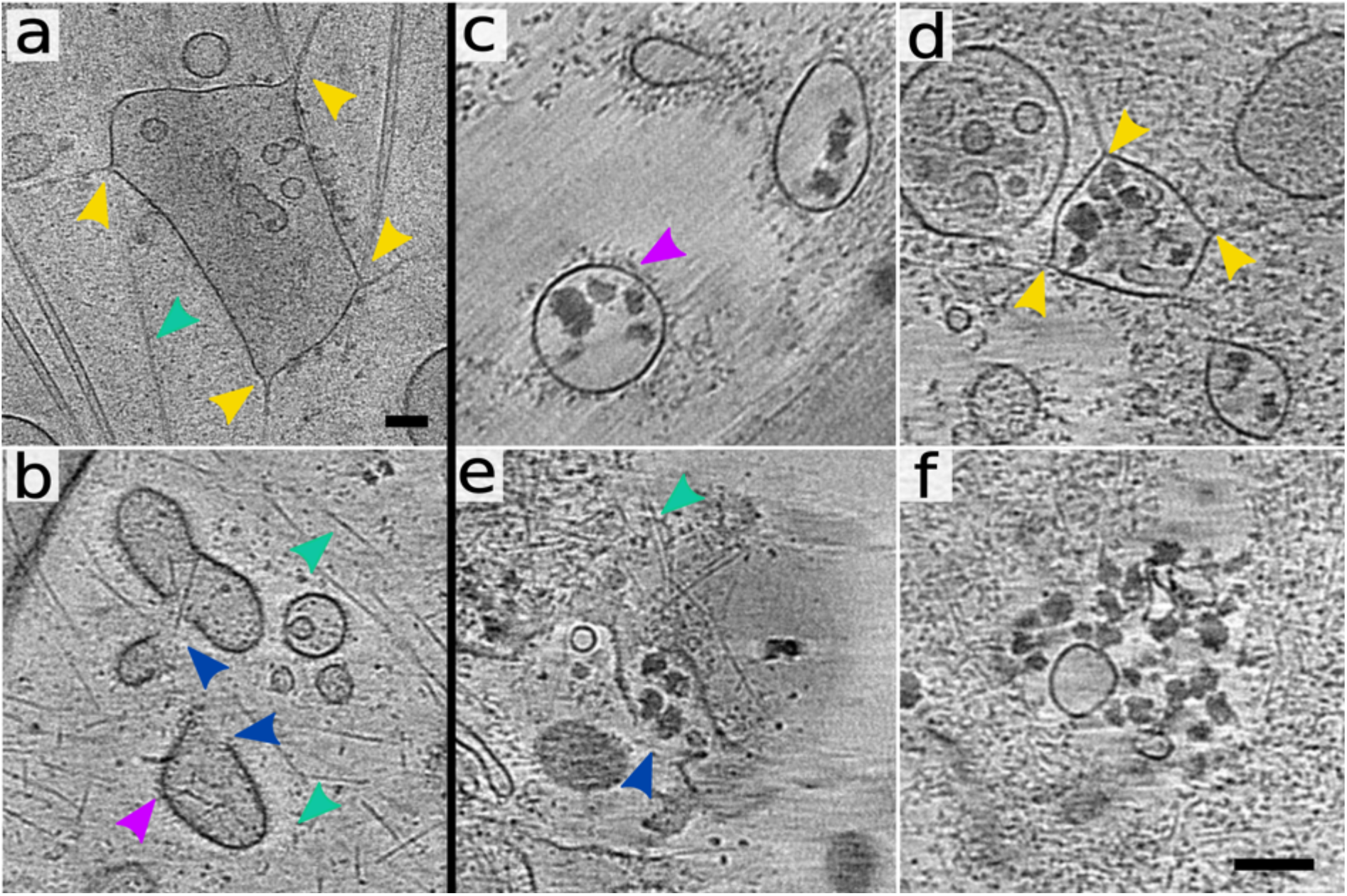
Endosome rupture in uninfected cells and release of VLDL from ruptured endosomes. The images show 1.66-nm-thick tomographic slices of cos-7 cells. **(ab)** Warped (a) and ruptured (b) endosomes in control cells. The positions of membrane warping are indicated by yellow arrowheads, and the openings are indicated by blue arrowheads. Coated area of an endosome is indicated by a magenta arrowhead. **(c-f)** Sections from cryo-tomograms of cos-7 cells sixty minutes after the addition of OsO_4_-stained VLDL complexes. **(b)** Single-membrane endosomes with smooth surfaces containing VLDL complexes. VLDL complexes are visible as electron-dense irregular objects. **(c)** Endosome with warped membrane forming cone-shaped protrusions. Selected tips of membrane warping are indicated by yellow arrowheads. Actin filaments attached to the tips of the membrane cones are visible. **(e)** Ruptured endosome releasing VLDL into the cytoplasm. The position of the opening in the endosome membrane is indicated by a blue arrowhead. **(f)** VLDL complexes in the cell cytoplasm. Scale bars 100 nm. Please note that panel (a) is shown at lower magnification than panels (b-f).

### Enteroviruses are released into cytoplasm from ruptured endosomes

The escape of enterovirus particles or genomes from endosomes is essential for the initiation of infection. Cells infected by rhinovirus 2, echovirus 18, echovirus 30, and enterovirus 71 contained ruptured endosomes with openings of sufficient size to enable diffusion of the viruses into the cytoplasm (Fig. 1a-b, 2g-j, S1d, S2d, S3d, Movie S1). Numerous enterovirus particles were in the vicinity of the disrupted vesicles. The studied viruses bind to different receptors at the cell surface and belong to distinct clades within the genus *Enterovirus*. Nevertheless, our results provide evidence that they all employ the same mechanism of endosome escape. Therefore, it is likely that other enteroviruses may use endosome rupture to reach the cytoplasm.

Endosome disruption was previously proposed as a genome delivery mechanism for major group rhinoviruses and rhinoviruses adapted to use heparan-sulfate as a receptor (*15, 46, 47*). In contrast, there is biochemical evidence that minor group rhinoviruses, including rhinovirus 2, release their genomes in late endosomes and translocate them into the cytoplasm through transmembrane pores (*35, 36*). These assumptions were based on *in vitro* analyses of the association of rhinovirus 2 with specific membrane fractions isolated from infected cells (*35, 36*). Our results indicate that at least some rhinovirus 2 particles reach cytoplasm via mechanism similar to that previously proposed for major group rhinoviruses.

Cryo-ET enables three-dimensional analysis of cell and virus structures; however, it does not allow monitoring of the progress of infection of individual virus particles over time. Therefore, it cannot be determined which, if any, of the observed particles would successfully initiate infection. Moreover, the enterovirus infectious unit is formed by about 1,000 particles (*67, 68*). To enable the visualization of enterovirus infection by cryo-ET, the cells were infected using a multiplicity of infection of 29 (please see Materials and methods for details). This high virus dose was required to enable the observation of multiple enterovirus particles in high-magnification tomograms, each of which captured only 0.02% of a cell volume. Therefore, our cryo-ET analyses do not provide proof that endosome disruption is the only enterovirus infection pathway. Nevertheless, we demonstrate that enteroviruses can be released from endosomes by membrane rupture. The involvement of membrane rupture in enterovirus infection under a low multiplicity of infection conditions is corroborated by the blockage of enterovirus infection by wiskostatin – an inhibitor of post-endocytic membrane remodeling, as is discussed below.

### Endosome rupture is triggered by a natural cellular mechanism

Lipid vesicles, including endosomes, are stable under physiological conditions, and their rupture requires a force that exceeds the forces holding the membrane together. The membrane warping and the shapes of the ruptured endosomes in enterovirus-infected cells indicate an external force pulling on the endosome membrane – a mechanism independent of the action of VP4 and the N-termini of VP1 subunits. Non-infected control cells contain a fraction of endosomes with ruptured membranes (Fig. 3b, S6), suggesting that endosome disruption can be triggered by a cellular mechanism without the direct involvement of virus components.

Rhinovirus 2 utilizes proteins from the family of LDLRs, including very low-density lipoprotein (VLDL) receptor, for cell binding and entry (*17*). We investigated VLDL cell entry to enable the comparison of the effects of the virus and native cargo on endocytosis mediated by the same receptor. The various stages of the development of VLDL-containing endosomes are similar to those of endosomes that contain rhinovirus 2 (Fig. 3b-f). Inside the endosomes, VLDL complexes detached from receptors due to acidic pH and lowered calcium ion concentration (*48–52*) and were distributed randomly (Fig. 3b-d). Membranes of some VLDL-containing endosomes warped (Fig. 3cd), and some of the tips of the membrane protrusions were attached to actin filaments (Fig. 3d). Finally, membranes of some VLDL-containing endosomes were ruptured (Fig. 3e), and VLDL diffused into the cytoplasm (Fig. 3f). Previous studies have shown that VLDL is transported to lysosomes for degradation and release of lipids and cholesterol to be incorporated into membranes (*69*). Our results do not exclude this canonical pathway; however, we show that a fraction of VLDL is released into the cytoplasm by endosome rupture via a mechanism morphologically similar to that utilized by enteroviruses.

### Blockage of post-endocytic membrane remodeling inhibits enterovirus infection

To test the hypothesis that endosome disruption, mediated by a cellular pathway, enables enteroviruses to reach the cytoplasm, we blocked post-endocytic membrane remodeling with wiskostatin. The binding of wiskostatin to N-WASP stabilizes the protein in autoinhibited conformation, prevents it from activating the Arp2/3 complex, and thus blocks processes of cellular membrane remodeling that depend on actin polymerization (*70, 71*). The infection of cells pre-treated with wiskostatin increased the odds of enterovirus genome release inside endosomes 10-fold relative to untreated cells (Fig. 4ae, Table S3). Most of the particles were trapped inside oblique endosomes; however, some were released into the cytoplasm (Fig. 4b). The inhibition of endosomal membrane remodeling by wiskostatin probably resulted in a prolonged retention of enterovirus particles in endosomes, where they released genomes. Relative to one-hour post-infection, echovirus 18 replicated its genome 20- and 35-fold, seven and nine hours post-infection, respectively (Fig. S7, Table S4). In contrast, in cos-7 cells pretreated with wiskostatin, the relative replication of echovirus 18 did not exceed 5-fold both seven and nine hours post-infection (Fig. S7). Thus, wiskostatin treatment restricts echovirus 18 infection relative to untreated cells. In summary, our results suggest that N-WASP-mediated actin polymerization is important for the disruption of endosomes, which enables the delivery of enteroviruses into the cytoplasm.

**Fig. 4.**
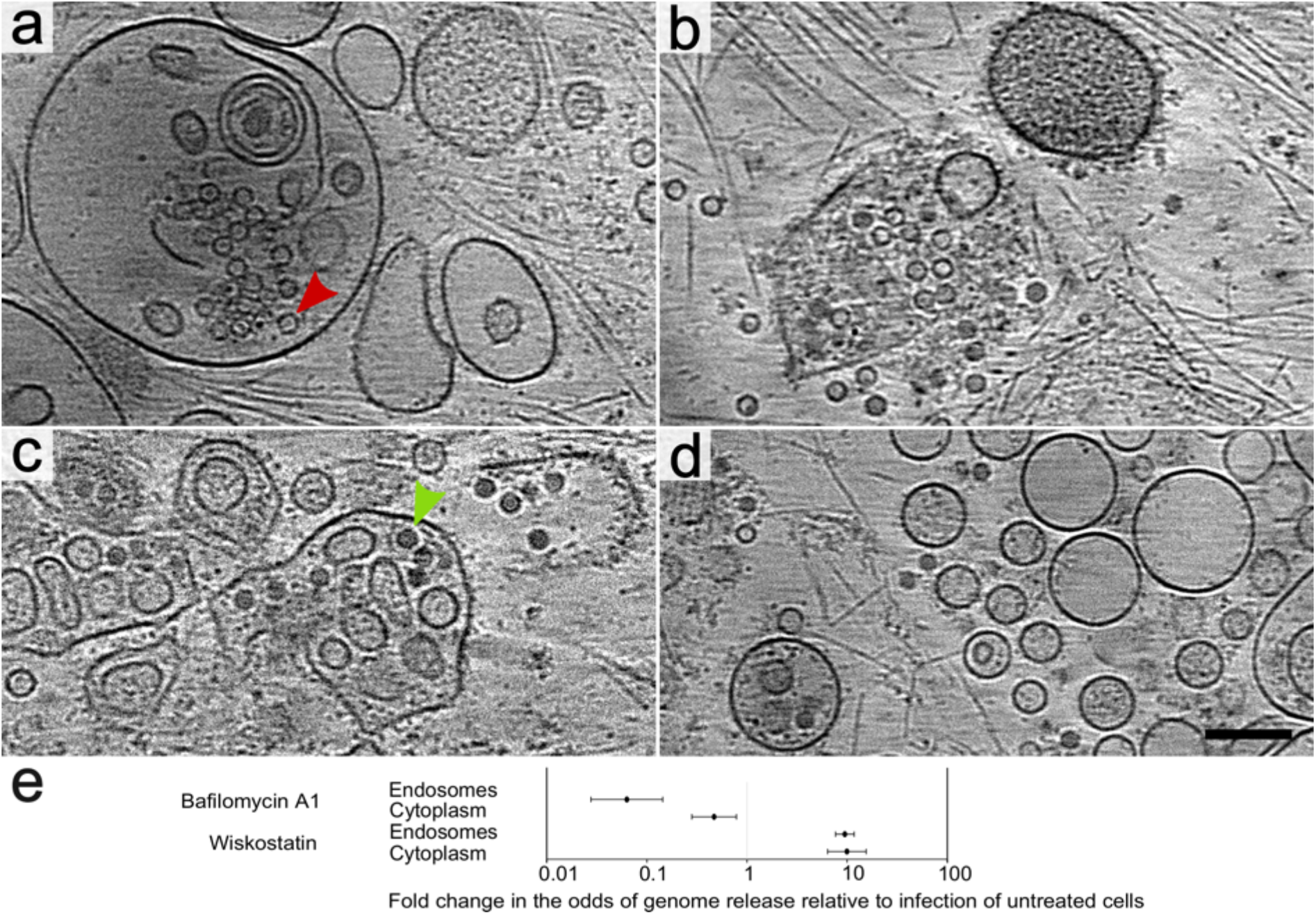
Effects of inhibitors interfering with endosome maturation on enterovirus cell entry. Panels (a-d) show 1.66-nm-thick tomographic slices from cos-7 cells pre-treated with the inhibitors for thirty minutes before infection and vitrified sixty minutes post rhinovirus 2-infection. **(ab)** Cells treated with wiskostatin, an inhibitor of N-WASP-mediated actin polymerization that is required for post-endocytic membrane remodeling. (a) Oblique endosomes containing empty particles of rhinovirus 2. A red arrowhead indicates a selected empty particle. (b) A rare occurrence of ruptured endosome in wiskostatin-treated cos-7 cells. **(cd)** Cells treated with bafilomycin A1, an inhibitor of endosome acidification. (c) Endosomes full of rhinovirus 2 virions. A green arrowhead indicates a selected virion. (d) Virions are released into the cytoplasm. Scale bar 100 nm. (e) Plot comparing differences in the odds of genome release from enterovirus particles in endosomes and cytoplasm of wiskostatin and bafilomycin A1-treated cells relative to the infection of untreated cells.

### Enteroviruses release genomes in endosomes and cytoplasm

Enterovirus genomes can be degraded by RNases, but it has been shown that the co-endocytosis of RNases with infecting viruses does not reduce enterovirus infectivity (*45, 72*). However, the mechanism that protects enterovirus genomes from degradation remains unknown. Here we show that both endosomes and the cytoplasm of infected cos-7 cells contained full and empty particles of the studied enteroviruses (Fig. 2, S1-4). In most cases, the cytoplasm contained a higher fraction of empty particles than endosomes (Fig. S4), which confirms previous findings that enteroviruses release their genomes in the endosomes and cytoplasm (*15*).

The studied enteroviruses differed in the speed of accumulation of empty particles in infected cells. The fastest to release its genome was echovirus 18, with 84% empty particles in the cytoplasm thirty minutes post-infection, and the slowest was enterovirus 71, with 2% empty particles sixty minutes post-infection (Fig. S4). The differences may be caused by (i) the viruses utilizing distinct receptors for cell entry, which may affect the speed of endocytosis and conditions in endosomes (*73*), (ii) different particle stability in acidic pH, which affects the timing of the initiation of genome release (*74*), and (iii) differences in the stability of empty particles of the enteroviruses, which may fall apart and thus would not be included in our statistics.

All the studied enteroviruses induce endosome disruption (Fig. 1, S1-3); therefore their virions and free genomes are released from endosomes into the cytoplasm. Nevertheless, the cell compartment where a genome is released probably influences the efficiency of infection, since the endosome lumen contains RNases that may cleave enterovirus genomes (*72*). The release of the genome in the cytoplasm may have a higher probability of successful initiation of infection, because it reduces the chances of genome degradation due to the dilution and putative deactivation of the RNases from endosomes in the cytoplasm. This may explain the previous observation that co-endocytosis of poliovirus and equine-rhinitis A virus with RNases did not prevent the virus infection (*45*).

### Activated particles are short-lived precursors of genome release *in vivo*

Cryo-tomograms of infected cells enable the straightforward identification of enterovirus particles and the determination of whether they contain genomes (Fig. 2, S1-3). However, distinguishing enterovirus virions from activated particles, both of which contain genomes, requires high-resolution information, since the two structures differ only in the details of their capsids (Table S1) (*26, 30, 75*). Single-particle reconstruction of rhinovirus 2 using electron micrographs collected on lamellipodia of infected cos-7 cells enabled the determination of the structures of virions and empty particles to resolutions of 4.7 and 7.1 Å, respectively (Fig. 1de, S8, Table S2). The classification procedures employed during the reconstruction of the genome-containing particles did not identify a sub-class of activated particles. Therefore, the infected cells probably only contained a small fraction of activated particles, which indicates that they are short-lived and rapidly converted to empty capsids. The fast release of genomes from activated particles *in vivo* contrasts with numerous previous *in vitro* studies in which the activated particles were sufficiently stable to enable the determination of their structures by cryo-EM or X-ray crystallography (*28, 30–32*). However, these experiments were performed at non-physiological pH, ion concentrations, and temperatures that may have stabilized the activated particles. Rapid genome release may be beneficial for enteroviruses *in vivo* because it accelerates the onset of virus translation and replication.

### Endosome acidification is not required for enterovirus escape from endosomes

Compounds that prevent the acidification of endosomes, including bafilomycin A1, block enterovirus infection to various extents (*24, 76–78*). *In vitro* experiments have demonstrated that acidic pH induces the formation of activated particles and genome release of many enteroviruses, which is prevented by bafilomycin A1 (*26–30, 32, 33*). Here we show that the treatment of cells with bafilomycin A1 leads to a 15-fold reduction in the odds of rhinovirus 2 genome release in the endosomes (Fig. 4ce). However, particles of rhinovirus 2 were released into the cytoplasm in bafilomycin A1-treated cells (Fig. 4d), and the odds of their genome release into the cytoplasm were only reduced 2-fold relative to the untreated cells (Fig. 4de). The observations that bafilomycin A1 treatment (i) does not prevent rhinovirus 2 escape from endosomes and (ii) has only a moderate effect on the probability of genome release in the cytoplasm, explain the limited effect of the compound on enterovirus infection described previously (*24, 76–78*). Furthermore, our results provide evidence that enterovirus escape from endosomes is independent of the activation of enterovirus particles and genome release.

### Summary

Based on previous results and our current analysis of enterovirus cell entry, we propose the following mechanism of enterovirus genome delivery: Enteroviruses enter cells by receptor-mediated endocytosis (Fig. 5) (*7–9, 15*). In endosomes, the studied enteroviruses detach from receptors (*31, 48–52*), and some of the particles release their genomes. N-WASP-mediated actin polymerization (*62, 63*) mediates the warping and rupture of enterovirus-containing endosomes (Fig. 5). Subsequently, enteroviruses diffuse from the ruptured endosomes to the cytoplasm, where additional particles may release their genomes. The cytoplasmic genome release may have a higher probability of resulting in productive infection than the endosomal one, because the RNases released from endosomes disperse and may be deactivated in the cell cytoplasm. Since the studied enteroviruses employ distinct receptors for cell entry but are all delivered into the cytoplasm by cell-mediated endosome disruption, we propose that most enteroviruses use endosome rupture to reach the cytoplasm.

**Fig. 5.**
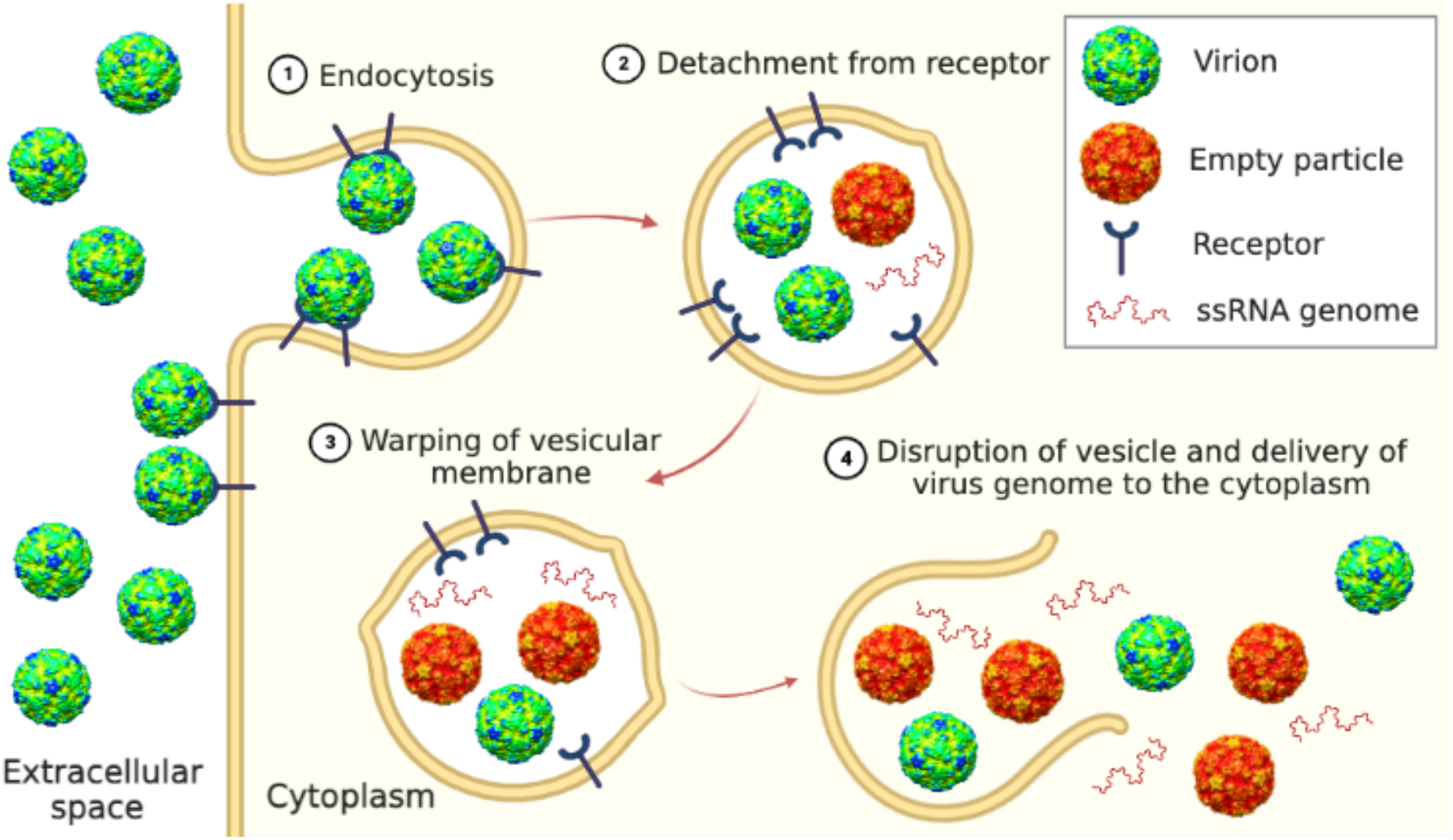
Overview of enterovirus cell entry. (1) Enterovirus virions bind to receptors at the cell surface and are endocytosed. (2) In endosomes, enteroviruses detach from receptors and are exposed to acidic pH. Some of the particles release their genomes. (3) Warping of membranes of virus-containing endosomes. (4) Rupture of endosome membrane, which depends on N-WASP-regulated actin polymerization, enables the diffusion of virions into the cytoplasm where additional particles release their genomes.

Our results identify endosome membrane remodeling as a new target for the development of therapeutics against enterovirus infections. Mutations in proteins involved in endocytosis (*79–82*), some of which are relatively common in the human population, affect the efficiency of endocytosis and endosome maturation. Individuals with mutated variants of the genes may have altered sensitivity to enterovirus infections, and their disposition may be a factor to consider when selecting a treatment strategy. Furthermore, endosome escape is a bottleneck for many therapeutics, because the carrier nanoparticles fail to exit endosomes (*83*). Targeting nanoparticles to bind receptors utilized by enteroviruses may improve the efficiency of nanoparticle endosome escape.

## Acknowledgments

We gratefully acknowledge the Cryo-electron Microscopy and Tomography, Biomolecular Interactions and Crystallization, and Proteomics core facility of CEITEC supported by MEYS CR (LM2018127). We wish to thank Alex J. Noble for helpful discussions.

## Author contributions

A.I. and P.Pl. designed research; A.I., L.M., Z.T., V.R., L.Š., J.M., K.Š., and J.N. performed research; A.I., L.M., Z.T., V.R., P.Pa., D.H., L.M., D.B., T.F., Š.V., J.N., and P.Pl. analyzed data; and A.I., L.M., Z.T., V.R., P.Pa., D.H., T.F., Š.V., and P.Pl. wrote the paper.

## Data availability

Cryo-EM electron density maps have been deposited in the Electron Microscopy Data Bank, https://www.ebi.ac.uk/pdbe/emdb/ under accession numbers EMD-15710 (rhinovirus 2 virion) and EMD-15711 (empty particle), and the fitted coordinates have been deposited in the Protein Data Bank, www.pdb.org under PDB ID codes 8AY4 and 8AY5, respectively. The authors declare that all other data supporting the findings of this study are available within the article and its Supplementary Information files, or are available from the authors upon request.

## Supplementary material for

### Supplementary materials and methods

#### Production and purification of enteroviruses

Rhinovirus 2 (strain HGP, ATCC-482) was propagated in HeLa (ATCC-CCL2) cells; echovirus 18 (strain METCALF, ATCC-VR-852), echovirus 30 (strain Bastianni, ATCC-VR-1660), and enterovirus 71 (strain MY104-9-SAR-97 (*87*)) were propagated in GMK cells (ATTC-CCL-81) cultivated in Dulbecco’s modified Eagle’s medium enriched with 10% fetal bovine serum (Sigma, F7534). For virus preparation, 50 tissue culture dishes with a diameter of 150 mm of cells grown to 100% confluence were infected with virus inoculum with a multiplicity of infection of 0.01 to 0.1. The infection was allowed to proceed for 24-72 hours, at which point more than 90% of the cells exhibited the cytopathic effect. Any remaining attached cells were removed from the dishes using cell scrapers, and the cell media were harvested. The cell suspension was centrifuged at 10,000 × g in a Beckman Coulter Allegra 25R centrifuge, TA-10-250 rotor at 10°C for 30 min. The resulting pellet was resuspended in 10 mL of phosphate-buffered saline (PBS) (10 mM Na_2_HPO_4_, 1.8 mM KH_2_PO_4_, 137 mM NaCl, and 2.7 mM KCl, pH 7.4, Sigma-Aldrich). The solution was subjected to three rounds of freeze-thawing by transfer between −80 and 37°C, and homogenized using a Dounce tissue grinder. Cell debris was separated from the supernatant by centrifugation at 3,100 × g in a Beckman Coulter Allegra 25R centrifuge, TS-5.1-500 rotor at 10°C for 10 min. The resulting supernatant was combined with media from the infected cells. Virus particles were precipitated by the addition of PEG-8000 and NaCl to final concentrations of 5% (w/v) and 0.5 M, respectively. The precipitation was allowed to proceed at 10°C with mild shaking (80 RPM) for 12 hours. The following day, the precipitate was pelleted at 10,000 × g in a Beckman Coulter Allegra 25R centrifuge, TA-10 rotor at 10 °C for 30 min. The supernatant was discarded, and the pelleted white precipitate was resuspended in 12 mL of PBS. MgCl_2_ was added to a final concentration of 5 mM, and the sample was subjected to DNAse (10 μg/mL final concentration) and RNAse (10 μg/mL final concentration) treatment for 30 min at ambient temperature. Subsequently, trypsin was added to a final concentration of 0.5 μg/mL, and the mixture was incubated at 37 °C for 10 min. pH 9.5 EDTA was added to a final concentration of 15 mM, and non-ionic detergent, NP-40^TM^ (Sigma Aldrich Inc.), was added to a final concentration of 1%. The mixture was spun down at 3100 × g in a Beckman Coulter Allegra 25R centrifuge, A-10 rotor at 10 °C for 10 min. Particles from the resulting supernatant were pelleted through a 30% (w/v) sucrose cushion in resuspension buffer (0.25 M HEPES, pH=7.5 and 0.25 M NaCl) by centrifugation at 210,000 × g in an Optima X 80 ultracentrifuge using a Beckman Coulter^TM^ type 50.2 Ti rotor at 10 °C for 2 h. The pellet was resuspended in 1.5 mL of PBS and loaded onto 60% (w/w) CsCl solution in PBS. The CsCl gradient was established by ultracentrifugation at 160,000 × g in an Optima X80 ultracentrifuge using a Beckman Coulter^TM^ SW41 Ti rotor at 10 °C for 18 h. The opaque bands containing the virus were extracted with a 20-gauge needle mounted on a 5 mL disposable syringe. The virus was transferred into PBS by multiple rounds of buffer exchange using a centrifugal filter device with a 100-kDa molecular weight cutoff.

#### Preparation of cryo-EM samples of cos-7 cells infected by enteroviruses

The 200 mesh holey carbon grids for transmission cryo-EM (Quantifoil, Au, R2/1) were sterilized with UV for 20 min and incubated in fetal bovine serum (Sigma) for 15 min to increase cell adhesion. The grids were placed into Thermo Scientific™ Nunc™ Lab-Tek™ Chamber Slide™ cell culture dishes, and 80 µL of cos-7 cell suspension (10^5^ cells / ml in Dulbecco’s modified Eagle’s medium enriched with 10% fetal bovine serum) were added to each well and cultivated for 16 h. Cells were washed twice with PBS and infected with about 5*10^10^ enterovirus particles in 5 µL of PBS (rhinovirus 2, echovirus 18, echovirus 30) or DMEM (enterovirus 71) per well. The infection was carried out at 37°C and 5% CO_2_ for 5 min. Subsequently the virus suspension was removed, fresh DMEM was added, and the cells were incubated at 37°C and 5% CO_2_ for the desired time. Not all virus particles from the inoculum attached and entered into the cells. Identification of enterovirus particles in tomograms of infected cells enabled calculation of the multiplicity of infection. The area of an average cos-7 cell was 2,500 μm^2^ (Fig. S9b) and the area of a reconstructed tomogram was 4,36 μm^2^. On average there were fifty enterovirus particles in a tomogram of an infected cell (Table S3, line infected cells un-treated by inhibitors). Therefore, an average cell was infected by 29,000 virus particles. The infectious unit to particle ratio of enteroviruses is about 1 to 10^3^ (*67, 68*). Therefore, the cells in our cryo-ET experiments were infected with a multiplicity of infection of 29. The high virus dose was required to enable observation of a representative number of enterovirus particles in the high-magnification tomograms, each of which represents only 0.17% of the area of an average cos-7 cell.

At the desired time post-infection, the cells were washed three times with PBS preheated to 37°C. Finally, 20 µL of PBS, containing 10 nm Gold tracers (Aurion, 210.133), preheated to 37°C were added to each well. PBS was removed by pipetting, and the grids were vitrified using a Vitrobot Mark IV. Cells were vitrified at 7-, 30-, 40- and 60-min post-infection. Control cells were vitrified using the same procedure, except for adding the virus.

#### Preparation of cryo-EM samples of cos-7 cells with VLDL

To enable the identification of very low-density lipoproteins (VLDL) in tomograms of cells, VLDL complexes were stained with osmium tetroxide using the following procedure: Twenty microliters of PBS containing VLDL at a concentration of 1.1 mg/ml (Calbiochem) were placed as a hanging drop into a 3.4 ml sealed container with 100 μl of 4% osmium tetroxide solution. The staining, using the vapor diffusion of osmium, was allowed to proceed for 8 min, during which the VLDL solution turned brown. Five microliters of osmium tetroxide-stained VLDL solution were added to cos-7 cells grown on 200 mesh holey carbon grids, as described above for the preparation of enterovirus-infected cells. The cells were incubated at 37°C for 5 min. Subsequently, the cells were washed three times with PBS preheated to 37°C and incubated in DMEM at 37° and 5% CO2 for 55 min. After the incubation, the cells were washed three times with PBS and vitrified, as described above.

#### Preparation of cryo-EM samples of cos-7 cells treated with wiskostatin or bafilomycin A1 and infected by rhinovirus 2

The protocols used to study the effects of wiskostatin and bafilomycin differed only in the concentration of the inhibitors, which were 50 µM for wiskostatin and 200 nM for bafilomycin. Cos-7 cells were grown on 200 mesh holey carbon grids, as described above for the preparation of enterovirus-infected cells. The cells were washed three times with PBS and then incubated in 5 µl of wiskostatin or bafilomycin dissolved in DMEM for 30 min. Subsequently, the medium covering the cells was replaced with 5 μl of DMEM containing rhinovirus 2 (5*10^10^ enterovirus particles) and wiskostatin or bafilomycin. After 5 min of incubation, the cells were washed with PBS and incubated in DMEM for another 55 min. After the incubation, the cells were washed three times with PBS and vitrified, as described above.

#### Calculating the effect of wiskostatin and bafilomycin A1 on the probability of enterovirus genome release

To determine the effect of wiskostatin and bafilomycin A1 on the probability that enterovirus particles release their genome, we calculated odds ratios (ORs) of full to empty particle conversion in drug-treated samples relative to an untreated control. Each odds ratio was calculated separately for each drug treatment and for particles in cytoplasm and in vesicles. For each odds ratio we performed statistical analysis and calculated 95% confidence intervals (95% CI), standard error (SE), standard normal deviate (z) and p-value (P). In wiskostatin-treated cells both in cytoplasm and in vesicles, the odds ratios showed a 10-fold increase in the probability of genome release relative to the control (cytoplasm: OR=9.93, 95% CI 6.38 - 15.46, SE=0.23, z=10.17, P<0.0001; vesicles: OR=9.48, 95% CI 7.65–11.74, SE=0.11, z=20.60, P<0.0001). In bafilomycin A1-treated samples, the odds ratio of genome release was reduced approximately two-fold in cytoplasm (OR=0.47, 95% CI 0.28–0.78, SE=0.26, z=2.90, P=0.0037) and about 15-fold in vesicles (OR=0.063, 95% CI 0.028–0.144, z=6.57, P<0.0001). Particle counts and the numbers of tomograms analyzed for each treatment are listed in Table S3.

#### Determining the effect of wiskostatin on enterovirus infection using reverse-transcription quantitative PCR (RT-qPCR)

Cos-7 cells were grown in 9.5 cm^2^ dishes to 80-90% confluency. Thirty minutes before infection, wiskostatin was added to the cell media to a final concentration of 50 µM. Subsequently, the cells were infected with echovirus 18 with a multiplicity of infection of 1-2. The cells were harvested 1, 7, and 9 hours post-infection. To isolate RNA, cells from one plate were lysed using 1 ml of TriPure reagent. The total RNA was isolated according to the manufacturer’s instructions (Roche). Total RNA was treated with 4 units of TurboDNase (AppliChem) followed by phenol : chloroform : isoamyl alcohol extraction and ethanol precipitation. The precipitated RNA was dissolved in RNAse-free water (Thermo Scientific). 1.5 μg of total RNA was used for cDNA preparation by reverse transcription (RT) with random hexamer primers and SuperScript III Reverse Transcriptase according to the manufacturer’s protocol (Invitrogen). The RT-qPCR was performed using a Roche LC 480 (Roche) RT-qPCR machine in FastStart SYBR Green Master 480 (Roche) with gene-specific primer pairs complementary to two different positions in echovirus 18 genome named V3 and V5, GAPDH, and macroglobulin (Table S4). Relative transcript abundance was calculated by the ∆∆Ct method (*88*). Data are displayed as mean values normalized to the housekeeping gene (GAPDH, microglobulin) and time point of 1 hour post-infection. The experiment was performed in three independent biological replicates. Results are shown as means and standard errors of the mean. p-values were calculated using a two-tailed Student’s *t*-test, and the error bars show the standard deviation with a p-value ***p<0,005.

#### Acquisition and analysis of cryo-electron tomography data

Tomographic tilt series of infected cells were collected using a ThermoFisher Titan Krios transmission electron microscope operating at 300 kV. The data were collected using a post-GIF Gatan K2 (Bioquantum 967) direct electron detector operating in zero-loss imaging mode with the width of the energy selecting slit set to 20 e^-^V. The grid was first screened to localize cells with thin lamellipodia suitable for cryo-ET data collection (Fig. S9). The tilt series were collected using SerialEM software (*89*) with a dose-symmetric tilt scheme (*90*) with an angular range of ±52° (2° increment) or ±60° (3° increment). With both settings, the total electron exposure was 50 e^-^/Å^2^. Individual images were saved as movies of 4 frames in counting mode. The data were collected at a magnification of 26,000 x, corresponding to a pixel size of 5.534 Å, and with defocus values ranging from −2.5 to −5.5 μm. Frames were aligned to compensate for drift and beam-induced motion during image acquisition using the program MotionCor2 (*91*). The resulting tilt series were aligned in IMOD using the patch tracking or fiducial marker-based alignment algorithms, binned three times, and the tomograms were calculated by weighted back-projection with a SIRT-like filter as implemented in the software package IMOD (*92*). Tomograms were denoised using the Topaz unet-3d-10a model (*93*).

#### Identification of the positions of virions, empty particles, ribosomes, and microtubules in tomograms

Aligned tilt series from IMOD were imported into an emClarity work environment, and the contrast transfer function (CTF) of the tilts was determined using emClarity (v 1.5.3.11). CTF-corrected tilt series were binned three times and reconstructed into tomograms (pixel size 16.57 Å). Template matching was performed by emClarity using resampled maps of ribosomes (EMD-8345), microtubules (EMD-12639), or HRV2 particles as templates. The set of template-matched particles was curated by inspecting each matched particle’s position in the tomogram. False-positive matches were removed. Positions and orientations of the matched particles were exported from emClarity into a homemade program (available at https://github.com/fuzikt/tomostarpy; placeback_subvolume.py), which was used to position the maps of the search templates into a volume that matched the size of the original tomogram. This resulted in separate maps of positioned maps of ribosomes, microtubules, and rhinovirus 2 virions and empty particles. The maps of the positioned components were overlaid in ChimeraX (*94*) together with volumes of manually segmented membranes, actin filaments, and the original tomogram.

#### Segmentation of cellular membranes and actin microfilaments in tomograms

The segmentation was performed using the software Amira (Thermo Fisher Scientific, version 2020.2). The membranes and filaments were manually segmented using the Brush tool and masking to select voxels belonging to membranes or microfilaments. The segmented labels representing membranes and microfilaments were then expanded using the feature “Grow selection in volume”. The labels were exported as three-dimensional maps in MRC format. Subsequently, the separate maps of membranes and microfilaments were overlaid with the volumes of virions, empty particles, ribosomes, and microtubules and the original tomogram using ChimeraX (UCSF ChimeraX, version 1.3 (2021-12-08)) (*94*). Pictures and movies were prepared using ChimeraX and the program FFMPEG (https://ffmpeg.org).

#### Acquisition and analysis of single-particle data

The data for single-particle analysis were collected using the same microscope and camera as with the cryo-ET data. Electron micrographs were recorded in the counting mode of the detector at a magnification of 105,000×, which corresponds to a pixel size of 1.34 Å. Total electron exposure was 50 e^-^ / Å^2^, and defocus values ranged from −1.8 to −3.0 μm. Movies of 40 frames were recorded for each 10 s of exposure. The frames from each exposure were aligned to compensate for drift and beam-induced motion using the program MotionCor2 (*91*). The resulting dose-weighted sums of aligned frames were used in the subsequent image processing steps, except for estimating contrast transfer function (CTF) parameters, which were determined from non-dose-weighted micrographs using the program gCTF v1.06 (*95, 96*).

Images of 2035 particles of rhinovirus 2 were manually boxed from micrographs of cells 30 and 40 min post-infection using cryolo_boxmanager from the software Cryolo 1.5.4 (*97*). The cellular localization of all of the boxed particles could not be determined with certainty due to limited fields of view and low contrast in the electron micrographs. Particles from within vesicles and cytoplasm were combined for the reconstruction. Particles were extracted with a box size of 448 pixels using RELION (*98*). 2D classification was employed to separate 1590 full and 332 empty particles. Initial models of full and empty particles with imposed icosahedral symmetry were independently generated *de novo* using Relion’s 3D initial model procedure. Subsequently, particles were subjected to several rounds of 3D classification using the initial models as references. Images of empty particles were binned to 224 pixels. Images of 1424 full particles and 332 empty particles were selected for reconstruction using the Relion 3D autorefine procedure. The resulting maps were used as initial models for subsequent 3D refinements. Icosahedral symmetry was imposed on the volumes during the refinement process. The final volumes were subjected to post-processing in RELION. The resolutions of the reconstructions were estimated according to the FSC 0.143 criterion.

To identify the putative sub-population of activated particles, all genome-containing particles were subjected to 3D classification using maps generated from PDB structures of the rhinovirus 2 virion (PDB: 1FPN) and activated particle (PDB: 4L3B) as starting models. Particles from the resulting classes were refined separately. In all cases the reconstructed maps corresponded to virions, or the resolution of the reconstructions was insufficient to distinguish whether the structure belonged to a virion or activated particle.

#### Comparison of maps for identification of particle type

The cryo-EM reconstruction of an empty particle of rhinovirus 2 was used for calibrating pixel size by comparing it to a map calculated from the PDB structure of an empty particle (PDB: 3TN9). The PDB structure was converted to a map using the software package XMIPP(*99*). The pixel size of the cryo-EM map of the empty particle was changed in small steps, and each time a cross-correlation coefficient comparing it to the map calculated from the PBD structure was calculated using the program Mapman (*100*). The highest cross-correlation was obtained for a pixel size of 1.324 Å. Previously determined PDB structures of the rhinovirus 2 virion (PDB: 1FPN)(*75*), activated particle (PDB: 4L3B)(*30*), and empty particle (PDB: 3TN9)(*26*) were converted to maps. The capsid cavity of cryo-EM reconstructions of the genome-containing particle was masked by applying a spherical mask with a diameter of 196 Å using the programs RELION 3.0.8 (*98*) and EMAN2 (*101*). The cryo-EM reconstructions and PDB-generated maps were low-pass filtered to a resolution of 4.7 Å using the program UCSF Chimera (*102*), and normalized using the program Mapman (*100*). Correlation coefficients comparing cryo-EM reconstructions and maps generated from PDB structures were calculated using the program Mapman (*100*).

#### Model building

The PBD structures of the virion (PDB: 1FPN) (*75*) and empty particle (PDB: 3TN9) (*26*) of rhinovirus 2 were rigid-body fitted into cryo-EM maps using the program Phenix (*103*). The structures were subjected to coordinate and B-factor refinement using the program Phenix (*103*).

### Supplementary figures

**Fig. S1.**
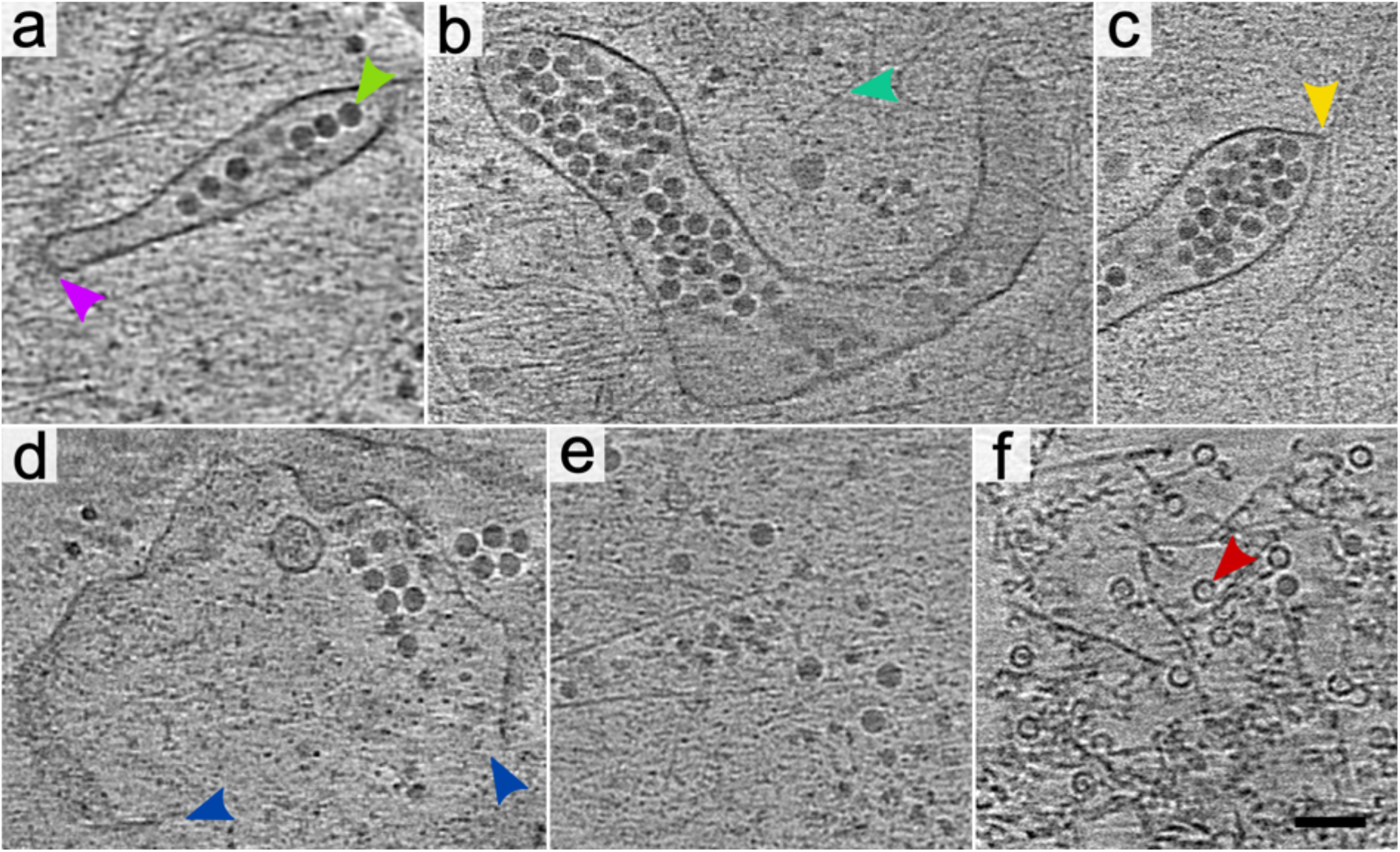
Release of echovirus 18 from endosomes into the cytoplasm. The images show 1.6 nm thick tomographic slices of echovirus 18-infected cos-7 cells 20-30 min post-infection. **(ab)** Endosomes containing echovirus 18 particles. The segment of the endosome covered with a protein coat in panel (a) is indicated by a magenta arrowhead. A selected virion is indicated by a green arrowhead. A selected actin microfilament in panel (b) is indicated by a cyan arrowhead. **(c)** Endosome with warped membrane forming a cone-shaped protrusion. The tip of the cone adjacent to the actin microfilament is indicated by a yellow arrowhead. **(d)** Ruptured endosome releasing echovirus 18 into the cytoplasm. The edges of the vesicle opening are indicated by blue arrowheads. **(ef)** Virions and empty particles of echovirus 18 in the cell cytoplasm. Scale bar 100 nm.

**Fig. S2.**
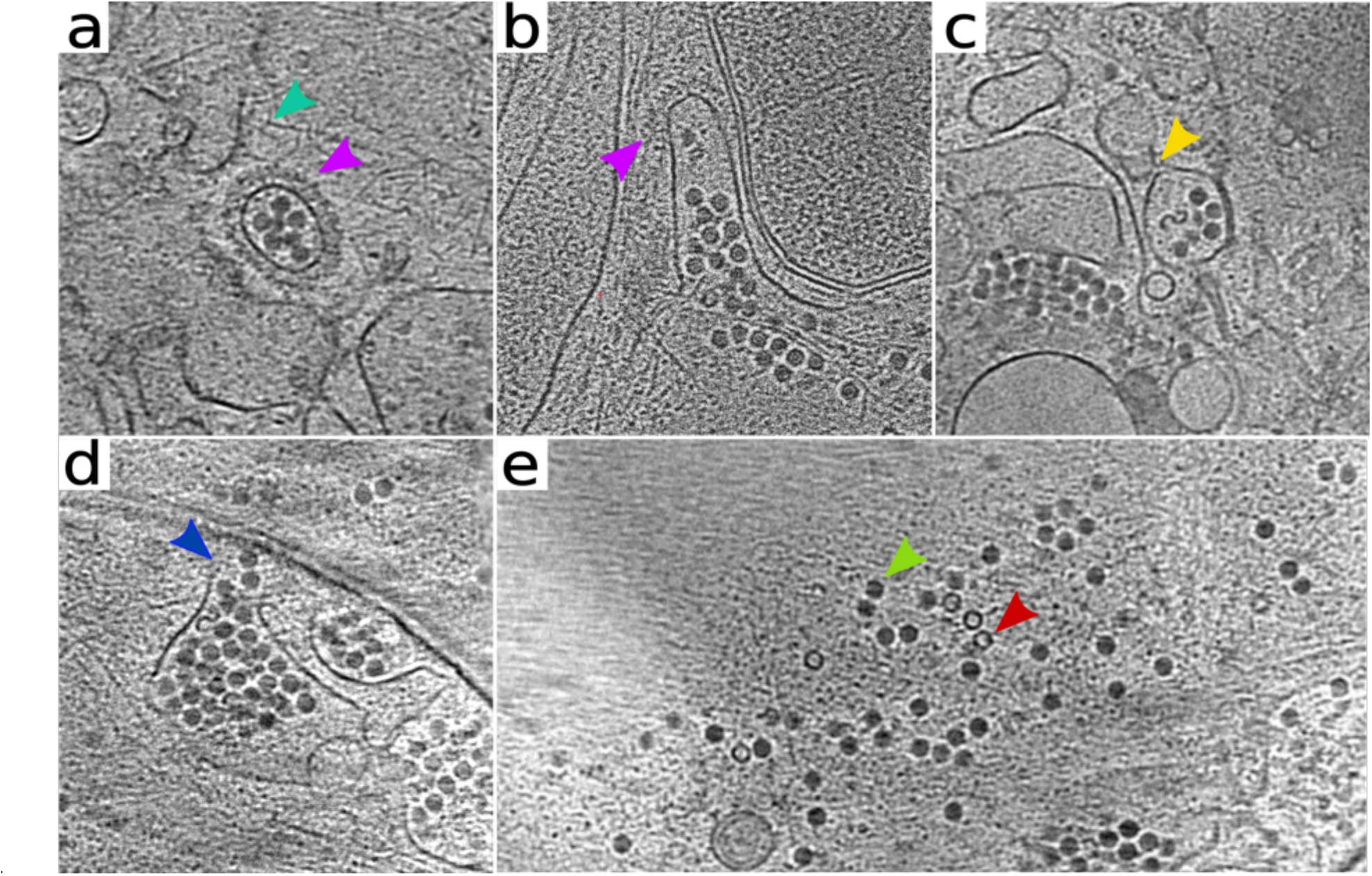
Release of echovirus 30 from endosomes into the cytoplasm. The images show 1.6-nm-thick tomographic slices of echovirus 30-infected cos-7 cells 30-60 min post-infection. **(a)** Endosome containing echovirus 30 particles. A segment of the endosome covered with a protein coat is indicated by magenta arrowhead. A selected actin microfilament is indicated by a cyan arrowhead. **(b)** Ruptured endosome with part of its membrane covered with a protein coat. The protein coat is indicated by a magenta arrowhead, a selected actin filament is indicated by a cyan arrowhead. **(c)** Endosome with warped membrane forming a cone-shaped protrusion. The tip of the cone is indicated by a yellow arrowhead. **(d)** Ruptured endosome releasing echovirus 30 into the cytoplasm. The opening in the endosome membrane is indicated by a blue arrowhead. The endosome is also warped and the tip of the membrane cone is indicated by a yellow arrowhead. **(e)** Virions and empty particles of echovirus 30 in the cell cytoplasm. Scale bar 100 nm.

**Fig. S3.**
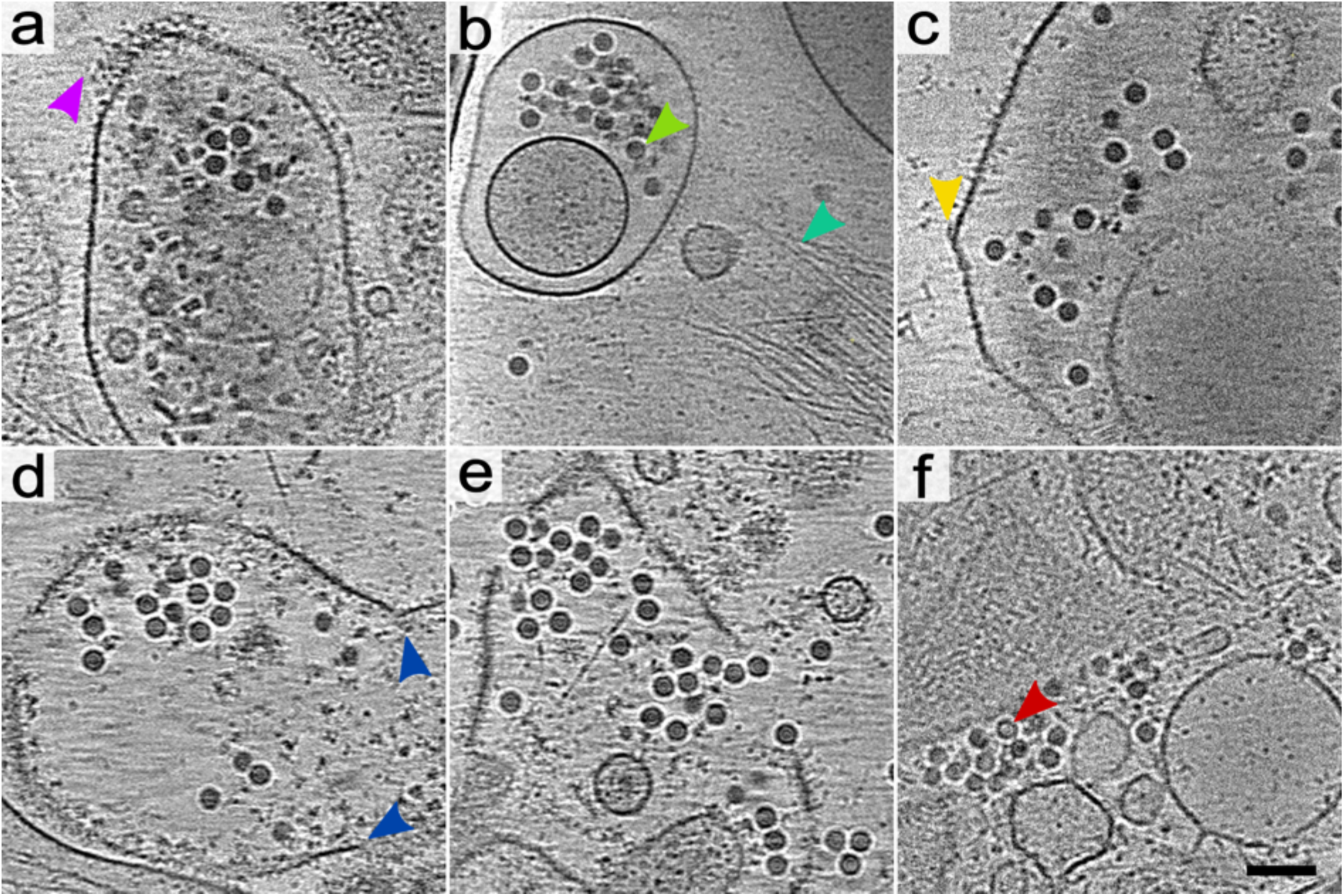
Release of enterovirus 71 from endosomes into the cytoplasm. The images show 1.6-nm-thick tomographic slices of enterovirus 71-infected cos-7 cells 30-60 min post-infection. **(a)** Endosome containing enterovirus 71 particles. A segment of the endosome covered with a protein coat is indicated by a magenta arrowhead. **(b)** Double-membrane endosome containing virus particles. Selected virion and actin microfilament are indicated by green and cyan arrowheads, respectively. **(c)** Endosome with warped membrane forming a cone-shaped protrusion. The tip of the cone is indicated by a yellow arrowhead. **(d)** Ruptured endosome releasing enterovirus 71 into the cytoplasm. The edges of the opening in the endosome membrane are indicated by blue arrowheads. **(ef)** Virions and empty particles in the cell cytoplasm. Selected empty particle in panel (f) is indicated by a red arrowhead. Scale bar 100 nm.

**Fig. S4.**
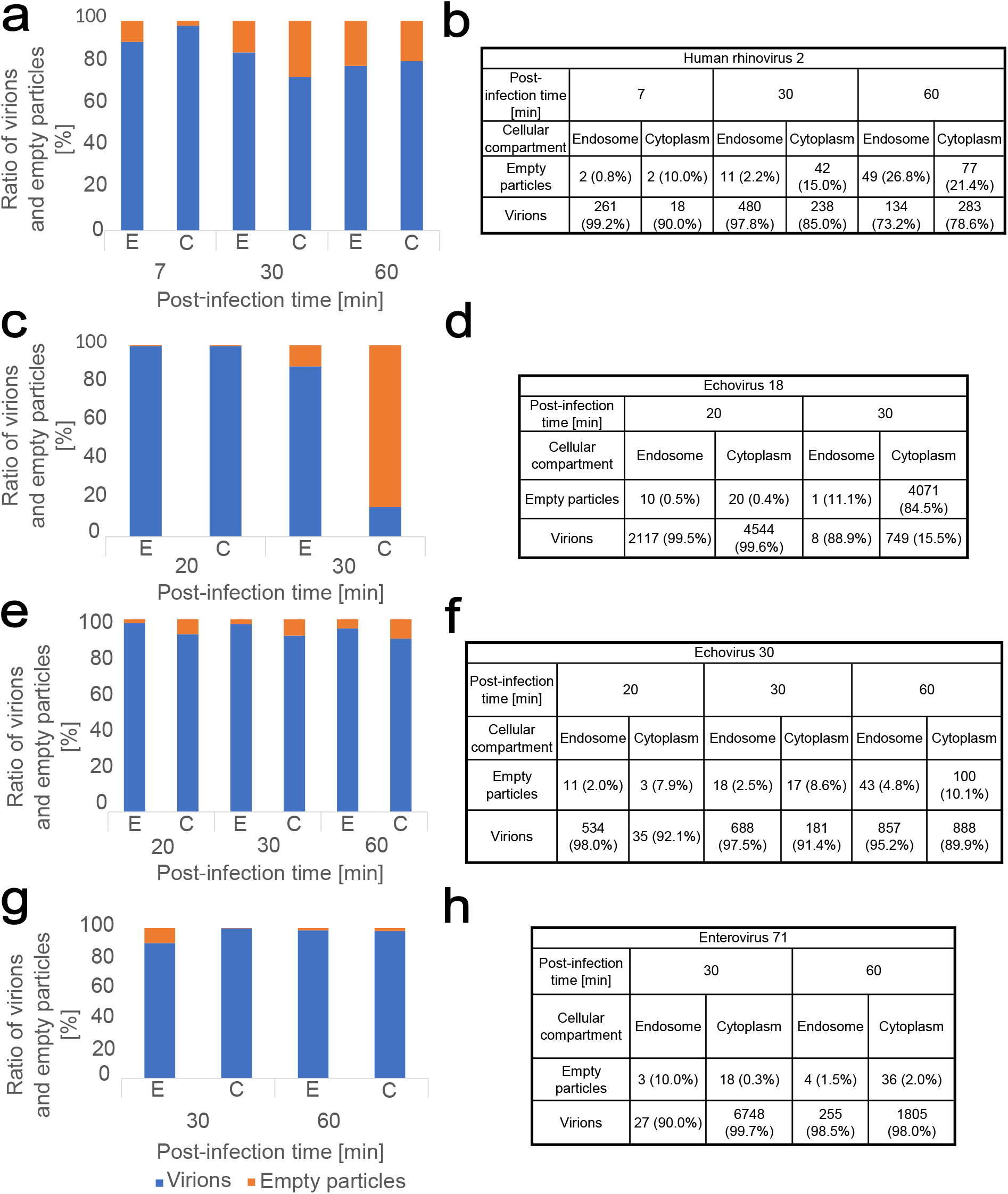
Fractions of virions and empty particles in endosomes and cytoplasm of infected cells. The graphs show the fractions of virions (blue) and empty particles (orange) in the cytoplasm and endosomes of infected cos-7 cells. Localizations of virus particles were determined using electron tomograms of cells. Only particles with certain localization were counted. The tables show absolute numbers of counted particles, the numbers in brackets indicate percentages of the particle types in each cellular compartment at a given time. Cells infected by **(ab)** rhinovirus 2, **(cd)** echovirus 18, **(ef)** echovirus 30, and **(gh)** enterovirus 71.

**Fig. S5.**
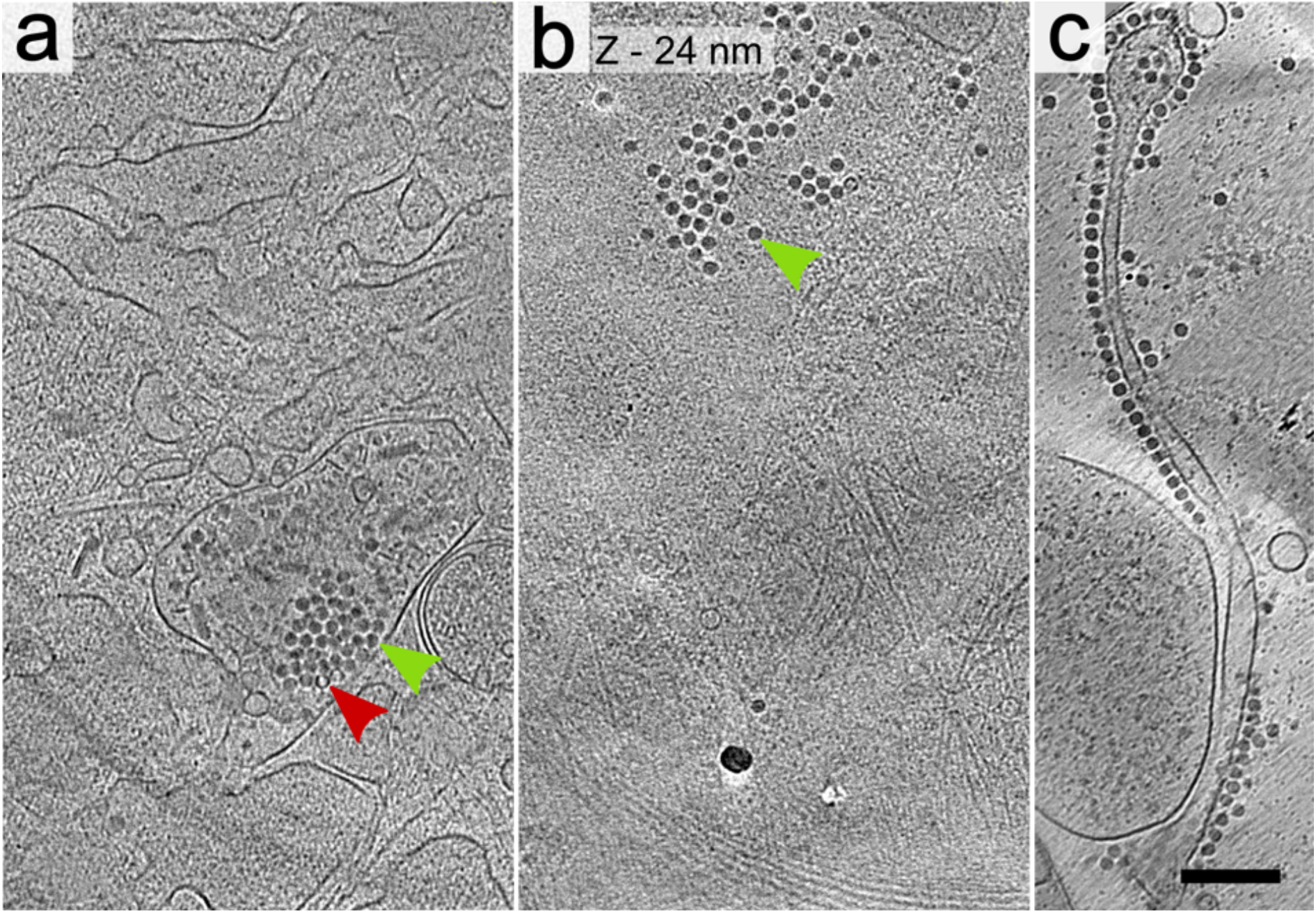
Attachment of enteroviruses to the cytoplasmatic membrane. The images show 1.6-nm-thick tomographic slices of cos-7 cells. **(ab)** Two tomographic slices of the same segment of echovirus 30-infected cos-7 cell showing (a) interior of the cell with virus particles in an endosome and (b) cell surface with attached virions. The green arrowheads indicate selected virions and the red arrowhead indicates a selected empty particle. **(c)** Filopodium of a cos-7 cell with virions of enterovirus 71 attached to its surface. Scale bar 200 nm.

**Fig. S6.**
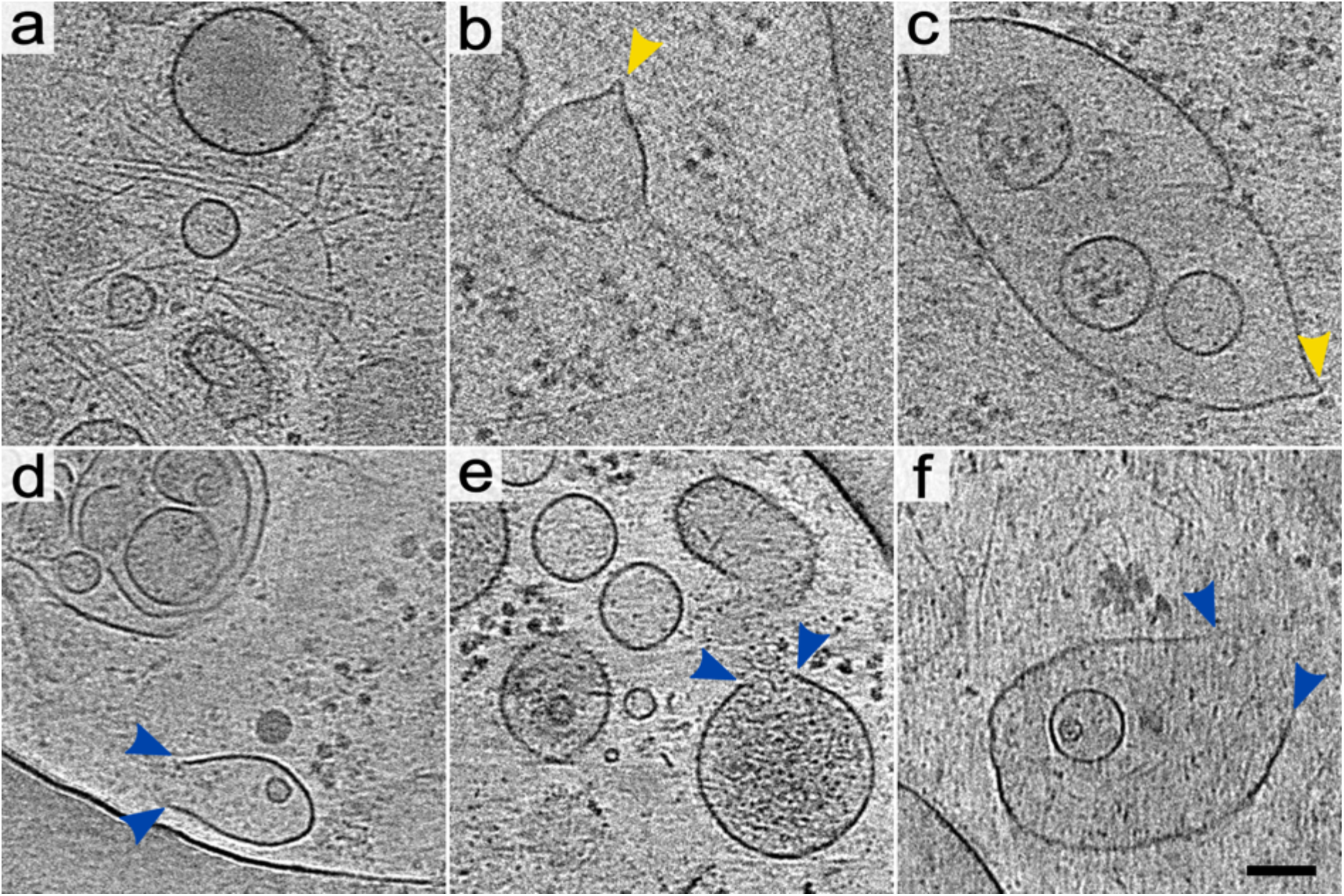
Warping and rupture of endosomes in control cells. The images show 1.6-nm-thick tomographic slices of cos-7 cells. **(a)** Endosomes with smooth membranes. **(bc)** Endosomes with warped membranes (tips of membrane cones are indicated by yellow arrowheads). **(d-f)** Disrupted endosomes (openings in the endosome membranes are indicated by blue arrowheads). Scale bar 100 nm.

**Fig. S7.**
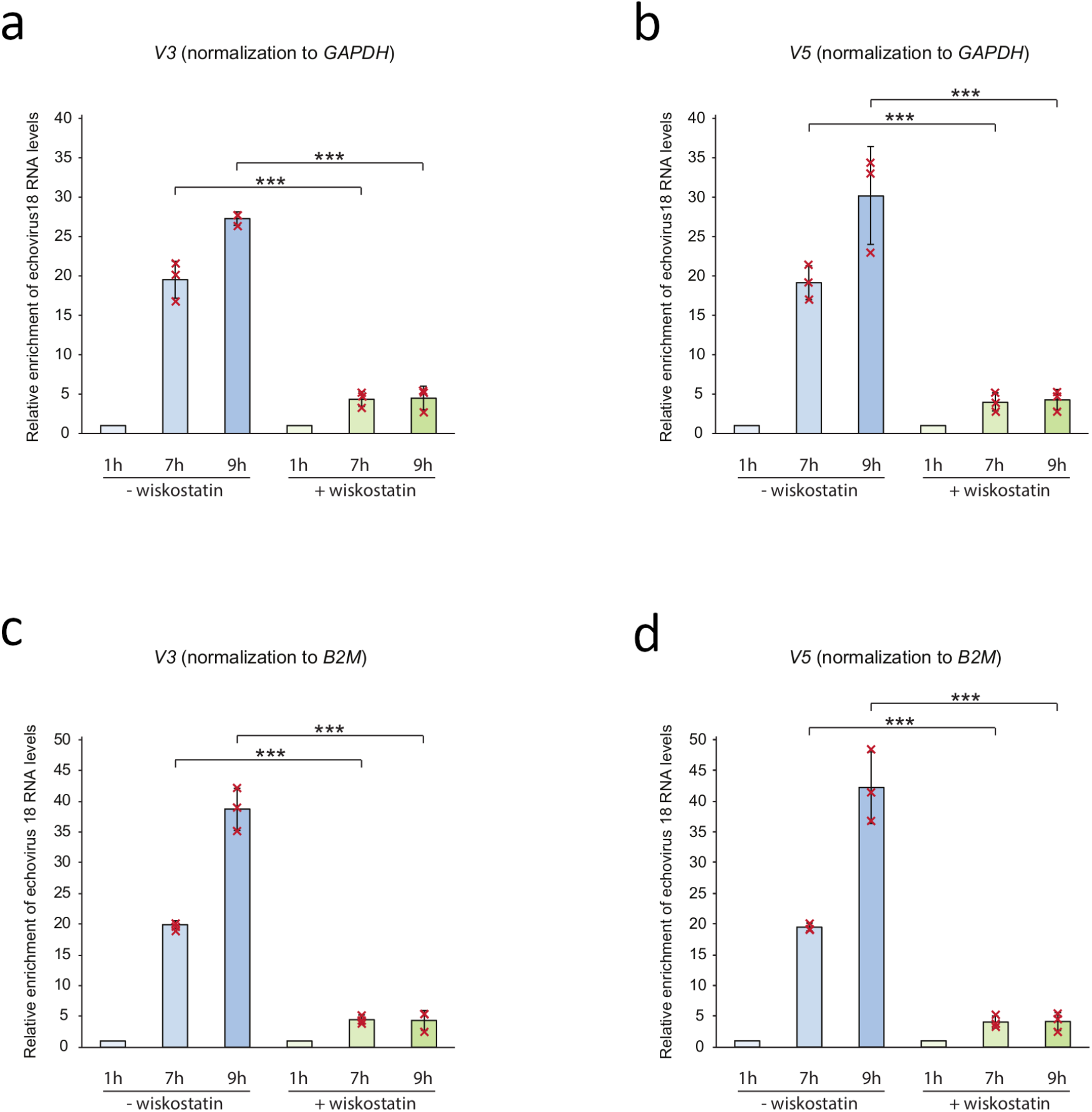
Wiskostatin inhibits infection of cos-7 cells by echovirus 18. RT-qPCR analysis of relative abundance of echovirus 18 RNA genomes in infected cos-7 cells. Relative levels of echovirus 18 genomes in control cells and cells treated with 50 µM wiskostatin 30 min before infection are shown at 1, 7, and 9 hours post-infection. The amounts of the virus genomes were quantified using V3 **(ac)** and V5 **(bd)** pairs of primers targeting distinct regions in echovirus 18 genome. The amounts of virus genomes were normalized by the ΔΔCt method to the house-keeping genes glyceraldehyde-3-phosphate dehydrogenase (ab) or ∆-2 microglobulin (cd) and to the time point of 1-hour post-infection. Standard deviations for three independent biological replicates are shown. *** indicates p<0,005. Primer sequences are listed in Table S4. Please see Materials and methods for details.

**Fig. S8.**
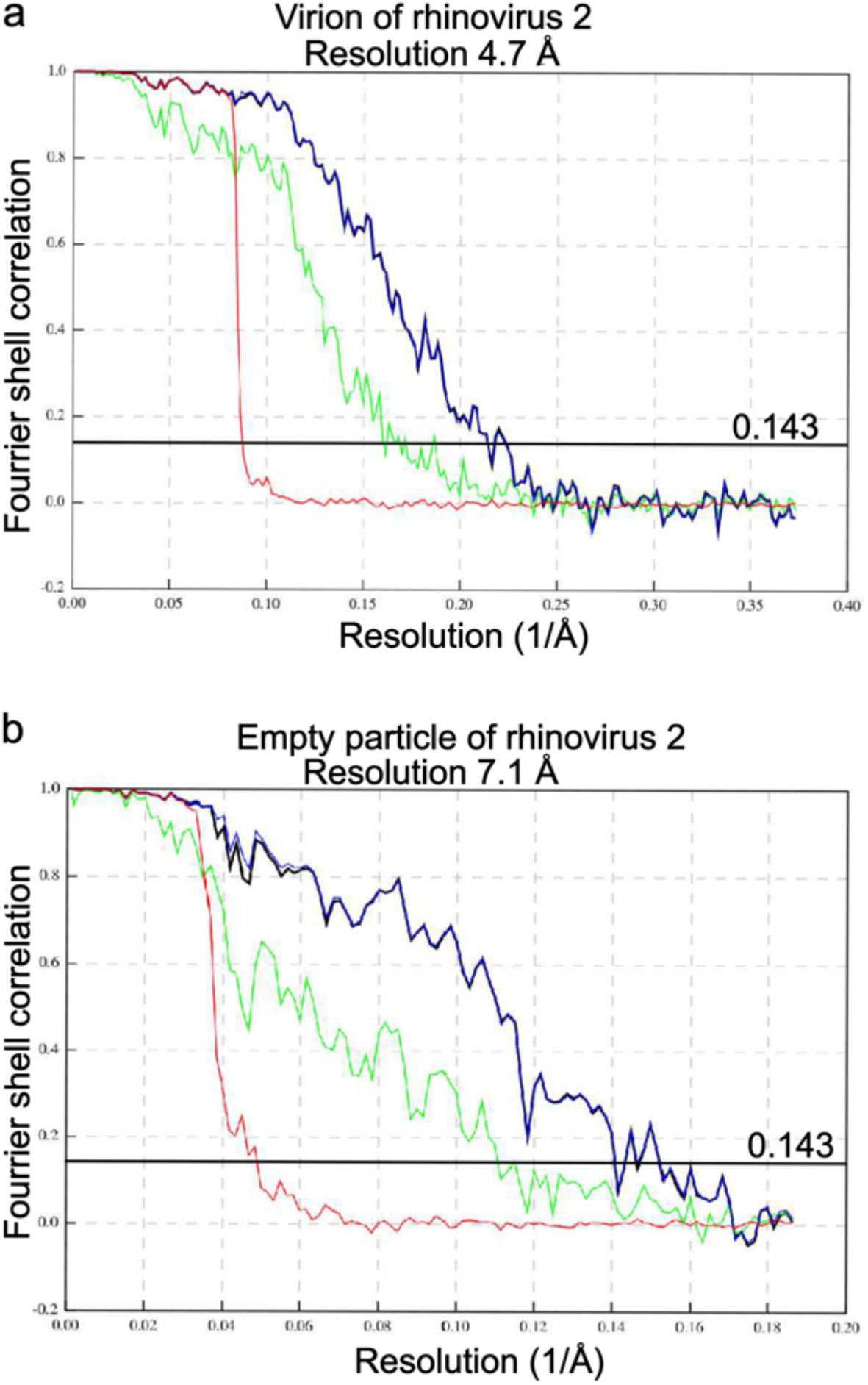
FSC curves of virion (a) and empty particle (b) of rhinovirus 2 reconstructed using micrographs recorded on infected cos-7 cells. FSC resolution threshold is 0.143.

**Fig. S9.**
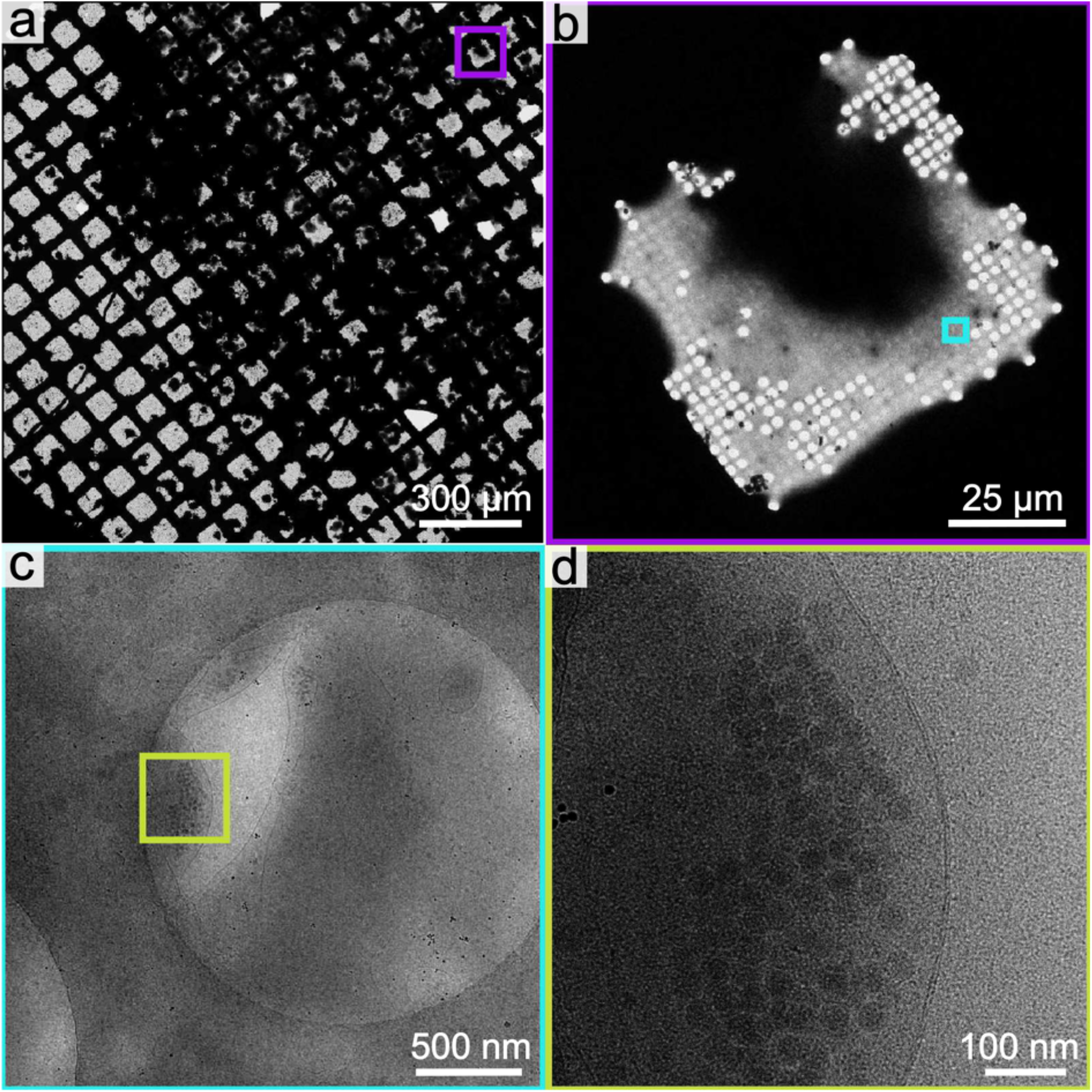
Series of images demonstrating the process of selection of an area for collecting tomographic tilt series of enterovirus-infected cos-7 cells. Panels (a-d) show a series of electron micrographs of cos-7 cells vitrified on a grid for electron microscopy recorded with increasing magnification. Every subsequent panel shows a magnified region indicated by a colored square in the previous one. **(a)** Overview of electron microscopy grid with cos-7 cells. **(b)** Image of a grid square with cos-7 cell. The dark area in the middle of the square represents a thick and electron non-transparent region of the cell. Grey areas represent cellular lamellipodia that are thin and electron-transparent. **(c)** Detail of cell lamellipodium with particles of rhinovirus 2 in vesicles. **(d)** High-magnification image of a segment of a vesicle containing rhinovirus 2 (d).

### Supplementary tables

**Table S1.** Cross-correlation coefficients comparing *in situ* cryo-EM reconstruction of genome-containing particle to maps calculated from structures of rhinovirus 2 virion, activated particle, and empty particle determined by X-ray crystallography.

|  | X-ray structures: |  |  |
| --- | --- | --- | --- |
|  | Virion | Activated particle | Empty particle |
| Cryo-EM reconstruction of genome-containing particle | 0.815 | 0.572 | 0.598 |
| X-ray structures: |  |  |  |
| Virion |  | 0.689 |  |
| Activated particle |  |  | 0.873 |

The cryo-EM reconstruction of the genome-containing particle was compared to maps calculated from PDB structures of the rhinovirus 2 virion (PDB code: 1FPN), activated particle (PDB code: 4L3B), and empty particle (PDB code: 3TN9) (26, 30, 75). Comparison of the maps calculated from PDB structures was used to determine the expected cross-correlation coefficients for capsids with similar structures (empty particle versus activated particle) and different structures (virion versus activated particle). For details, please see Supplementary materials and methods.

**Table S2.** Cryo-EM data and structure quality indicators.

| Parameters | Rhinovirus 2 |  |
| --- | --- | --- |
|  | Virion | Empty particle |
| Detector | Bioquantum K2 | Bioquantum K2 |
| Detector acquisition mode | counting | counting |
| Nominal Magnification | 105,000 | 105,000 |
| Voltage (kV) | 300 | 300 |
| Electron exposure on a sample (e<sup>-</sup>/Å<sup>2</sup>) | 50 | 50 |
| Target defocus range (μm) | -1.8 to -3.0 | -1.8 to -3.0 |
| Calibrated pixel size (Å) | 1.324 | 1.324 |
| Symmetry imposed | I4 | I4 |
| Number of collected movies | 2,468 | 2,468 |
| Fractions per movie | 40 | 40 |
| Number of initial particle images | 2,035 | 2,035 |
| Number of particle images used in recon. | 1,424 | 332 |
| Initial model | *de novo* | *de novo* |
| Map resolution (Å) at FSC=0.143 | 4.7 | 7.1 |
| **Refinement and validation statistics** |  |  |
| *Model composition* |  |  |
| Non-hydrogen atoms (one protomer) | 12,129 | 11,499 |
| Protein residues (one protomer) | 781 | 738 |
| *B-factor* |  |  |
| Protein (Å<sup>2</sup>) | 171.12 | 168.27 |
| *RMSD* |  |  |
| Bond lengths (Å) | 0.012 | 0.012 |
| Bond angles (°) | 1.964 | 1.972 |
| *Validation* |  |  |
| MolProbity score | 2.33 (99th percentile) | 2.57 (98th percentile) |
| ClashScore | 7.52 (97th percentile) | 13.42 (97th percentile) |
| *Rotamers* |  |  |
| Favored (%) | 87.22 | 86.75 |
| Poor (%) | 3.21 | 3.29 |
| *Ramachandran plot* |  |  |
| Favored (%) | 90.09 | 90.18 |
| Poor (%) | 0.65 | 0.68 |
| *Database entry* |  |  |
| EMDB | 15710 | 15711 |
| PDB | 8AY4 | 8AY5 |

**Table S3.** Particle counts, and the numbers of tomograms used to determine the effects of wiskostatin and bafilomycin on the probability of genome release from individual particles.

|  | Virions in cytoplasm | Empty particles in cytoplasm | Virions in vesicles | Empty particles in vesicles | Total particles | Number of analyzed tomograms |
| --- | --- | --- | --- | --- | --- | --- |
| Infected untreated cells | 233 | 34 | 659 | 156 | 1082 | 23 |
| Infected wiskostatin-treated cells | 98 | 142 | 362 | 812 | 1414 | 16 |
| Infected bafilomycin A1-treated cells | 453 | 31 | 402 | 6 | 892 | 11 |

**Table S4.**
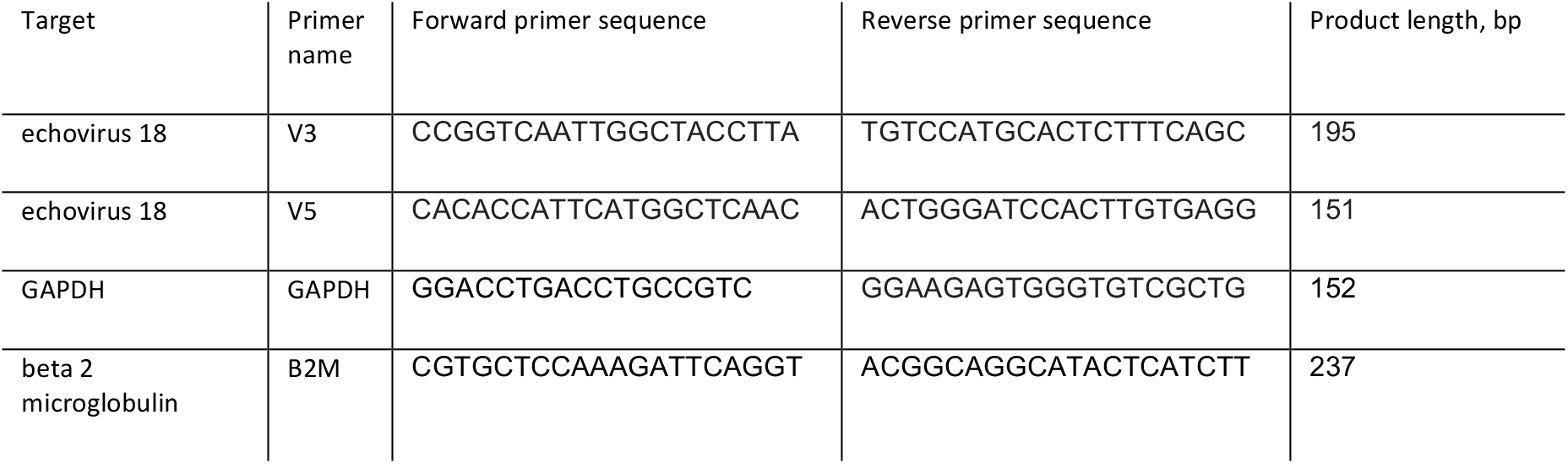
Primers used for RT-qPCR quantification of the effect of wiskostatin on enterovirus infection.

### Supplementary movie legends

**Movie S1: Cytoplasm of rhinovirus 2-infected cell imaged using cryo-ET.** The movie shows a tomographic reconstruction of an infected cos-7 cell 60 min post-infection. The positions of virions, empty particles, ribosomes, and microfilaments were identified using template matching as implemented in EMclarity (*86*), and the corresponding high-resolution structures were positioned into the tomogram. Microfilaments and membranes were segmented manually.

**Movie S2. Attachment of echovirus 30 to cytoplasmatic membrane.** The movie shows tomographic slices of infected cos-7 cell 30 min post-infection. Virions of echovirus 30 are attached to the cell surface, whereas other particles are in an endosome inside the cell.

